# Autosomal dominant Retinitis Pigmentosa caused by the rhodopsin isoleucine 255 deletion features rapid neuroretinal degeneration, decreased synaptic connectivity, and neuroinflammation

**DOI:** 10.1101/2024.08.29.610258

**Authors:** Bowen Cao, Yu Zhu, Alexander Günter, Ellen Kilger, Sylvia Bolz, Christine Henes, Regine Mühlfriedel, Mathias W. Seeliger, François Paquet-Durand, Blanca Arango-Gonzalez, Marius Ueffing

## Abstract

Retinitis Pigmentosa (RP) is a group of inherited retinal diseases that initially affects rod photoreceptors and causes progressive vision loss and blindness. Mutations in rhodopsin (*RHO*) can cause both autosomal recessive (ar) and dominant (ad) forms of RP, yet, the underlying degenerative mechanisms remain largely unknown, rendering the disease untreatable. Here, we focus on an in-frame, 3-base pair deletion, eliminating the isoleucine residue at codon 255 (*i.e., RHO*^ΔI255^) and resulting in adRP.

We generated a novel knock-in mouse homologous to the human *RHO*^ΔI255^ mutation. This new mouse model displays a severe disruption of photoreceptor structure and function, as is seen in human patients. Our results indicate that this form of RP is a systems disease of the neuroretina that also impacts neuronal connectivity of bipolar- and horizontal cells, initiates neuroinflammation, and reduces the structural and functional integrity of the retina.

Typical for adRP, *Rho*^ΔI255^ mice exhibit primary rod photoreceptor loss, followed by secondary cone degeneration, rhodopsin protein (RHO) mislocalization, progressive shortening of outer segments (OS), and disorganized OS structures. Subsequently, increasing gliosis, morphologic abnormalities of the inner retina, and impaired cone-driven visual function developed. In adRP, a single mutated allele is sufficient to cause the disease, as confirmed here in *Rho*^ΔI255/+^ heterozygous animals, where most photoreceptors were lost within two months after birth. Compared to this, homozygous *Rho*^ΔI255/ΔI255^ mutants exhibit an accelerated onset and even faster progression of retinal degeneration. The degeneration of *Rho*^ΔI255^-mutant photoreceptors was linked to the activation of both caspase- and calpain-type proteases, as well as poly(ADP-ribose) polymerase (PARP), indicating a parallel execution of both apoptotic and non-apoptotic processes.

In conclusion, our data indicate that this form of RP affects the neuroretina beyond photoreceptor cell loss sharing features typical for other degenerative central nervous systems diseases, an insight, which may bear critical impact to understand and eventually develop treatment for these currently untreatable forms of blindness.

**Author summary:** Dominant mutations in the human rhodopsin gene are among the most common causes for the blinding disease retinitis pigmentosa (RP). To date, the underlying pathophysiological mechanisms are still largely unknown and dominant RP remains untreatable. Here, we introduce a new knock-in mouse model carrying the dominant human *Rho*^Δ*I255*^ mutation. As in humans, the *Rho*^Δ*I255*^ mouse suffers from a rapid degeneration of rod photoreceptors followed by subsequent cell death of cone photoreceptors and complete loss of visual function. The new mouse model displays sign of neuroinflammation and the concomitant activation of both apoptotic and non-apoptotic cell death mechanisms. These results will likely stimulate further studies into the degenerative processes governing dominant RP and may facilitate future therapy development for inherited retinal diseases that are still untreatable to this day.

## Introduction

Neurodegenerative diseases are governed by a multiplicity of pathological alterations that eventually lead to neuronal dysfunction and cell death. Retinal degenerations are specific forms of neurodegeneration in the eye, primarily affecting photoreceptors. Among these, Retinitis pigmentosa (RP) denotes a group of hereditary retinal degenerations that primarily affects rod photoreceptor cells and leads to a progressive degeneration of the outer retina. RP exhibits a high degree of genetic heterogeneity, with over 80 different genes implicated in its pathogenesis, including both recessive and dominant mutations. The primary rod degeneration is followed by a secondary degeneration of cone photoreceptors, eventually culminating in complete blindness [1]. Rhodopsin (RHO) is a prototypical G protein-coupled receptor (GPCR) essential to rod photoreceptors. Over 150 distinct mutations in the human *RHO* gene are associated with 25% of autosomal dominant RP cases (adRP) [2]. Most of these mutations lead to defects in the biosynthesis and trafficking of the RHO protein and, ultimately, to the death of photoreceptors [3, 4].

RHO is synthesized in the rod inner segment and trafficked to the disc membrane of the rod outer segment via the connecting cilium. RHO is the most abundant protein in rod outer segments, representing approximately 90% of all proteins in this compartment [5]. The high protein expression of about six million molecules per rod and day, and the continuous renewal of the rod outer segments place a considerable metabolic burden on photoreceptors [6]. The biosynthesis of RHO requires highly coordinated regulation of *RHO* gene expression, protein translation and folding, glycosylation, palmitoylation, and transport to the OS. Dysregulation of any of these steps can lead to rod stress and death [7]. This may explain why even seemingly minor alterations to the *RHO* protein can have deleterious effects on rod outer segment structure and rod viability.

The first deletion mutation described in the human *RHO* gene involves a 3-bp deletion, eliminating one of two isoleucine residues at codons 255 and 256 within the proteińs sixth transmembrane domain [8]. The resulting mutant *RHO* allele is termed *RHO*^ΔI255^. The initial identification of this mutation emerged within a British pedigree [8] and was subsequently found also in Belgian [9], German [10], Chinese [11], and Korean families [12]. Its widespread geographical distribution emphasizes the importance of this gene defect and the need for further investigation. Patients carrying this mutation exhibit pronounced RP degeneration features, including night blindness from childhood and progressive vision loss over their lifetime [8, 13]. Despite its discovery three decades ago, the pathogenic mechanisms associated with the *RHO*^ΔI255^ mutation and adRP remain elusive. The deletion of isoleucine 255 has been categorized as a class II *RHO* mutation, characterized by the presence of RHO aggregates, full or partial retention of misfolded RHO within the endoplasmic reticulum (ER), and compromised RHO pigment production [14–16]. Our initial studies in HEK293 and COS-7 cells indicate that the mutant protein induces aggregate formation in the ER, recruiting the wild-type RHO into the aggregate pool and leading to degradation via the ER-associated degradation (ERAD) pathway [17].

To overcome the limitations of *in vitro* RP models, *in vivo* models are needed and so already in the late 1990s a transgenic mouse model carrying multiple copies of the *Rho*^ΔI255^ allele was generated. However, these transgenic animals displayed a very dramatic phenotype and from a genetic standpoint cannot be considered an exact model for autosomal dominant RP, since the mutation was not inserted in a site-specific manner [18]. To create an improved model that represents the situation in patients as closely as possible, we generated an *in vivo Rho*^ΔI255^ knock-in mouse model. This mouse model was then used to comprehensively investigate the pathophysiology and cell death pathways associated with this mutation. The phenotype of the novel *Rho*^ΔI255^ mouse line was strikingly similar to that observed in humans, with progressive rod and cone photoreceptor cell death and a decline in visual function. This was accompanied by RHO mislocalization, activation of glial cells, and loss of retinal structural integrity. In addition, the *Rho*^ΔI255^ mutation-induced degeneration triggered both classical apoptotic and alternative, non-apoptotic mechanisms of photoreceptor cell death [19]. Consequently, the new mouse model bears the potential to investigate pathologic mechanisms and to develop pre-clinical concepts for future therapeutic interventions.

## Results

### 1. *Rho*^ΔI255^ causes rapid retinal degeneration in mice with marked regional differences

Homologous recombination in mouse embryonic stem cells was employed to create the *Rho*^ΔI255^ knock-in mouse model [20], with C57BL/6J mice serving as the genetic background. This background was selected to avoid the introduction of the *rd8* mutation present in some other C57BL/6 mouse lines [21]. The embryonic stem cells were injected into mouse blastocysts, and after confirmation of germ-line transmission in the resultant chimeric animals, these were bred to homozygosity. Homozygous *Rho*^ΔI255/ΔI255^ mice were crossbred with C57BL/6J *Rho*^+/+^ animals (wild-type: WT) to obtain heterozygous *Rho*^ΔI255/+^ mutants for subsequent investigations.

We first verified whether mutant animals show a photoreceptor degeneration comparable to the human disease condition. Analyzing the retina after enucleation of the eyes, we employed the TUNEL assay at different postnatal (PN) days (Fig. 1A, Fig. S1A). A quantitative analysis of the percentage of dying TUNEL-positive cells in the outer nuclear layer (ONL) of both *Rho*^ΔI255/ΔI255^ and *Rho*^ΔI255/+^ retinae revealed increased photoreceptor cell death compared to WT. Homozygous *Rho*^ΔI255/ΔI255^ mice exhibited an early peak of photoreceptor degeneration at PN18, whereas heterozygous *Rho*^ΔI255/+^ mice displayed a later peak in retinal cell death at PN20 (Fig. 1B).

**Figure 1.**
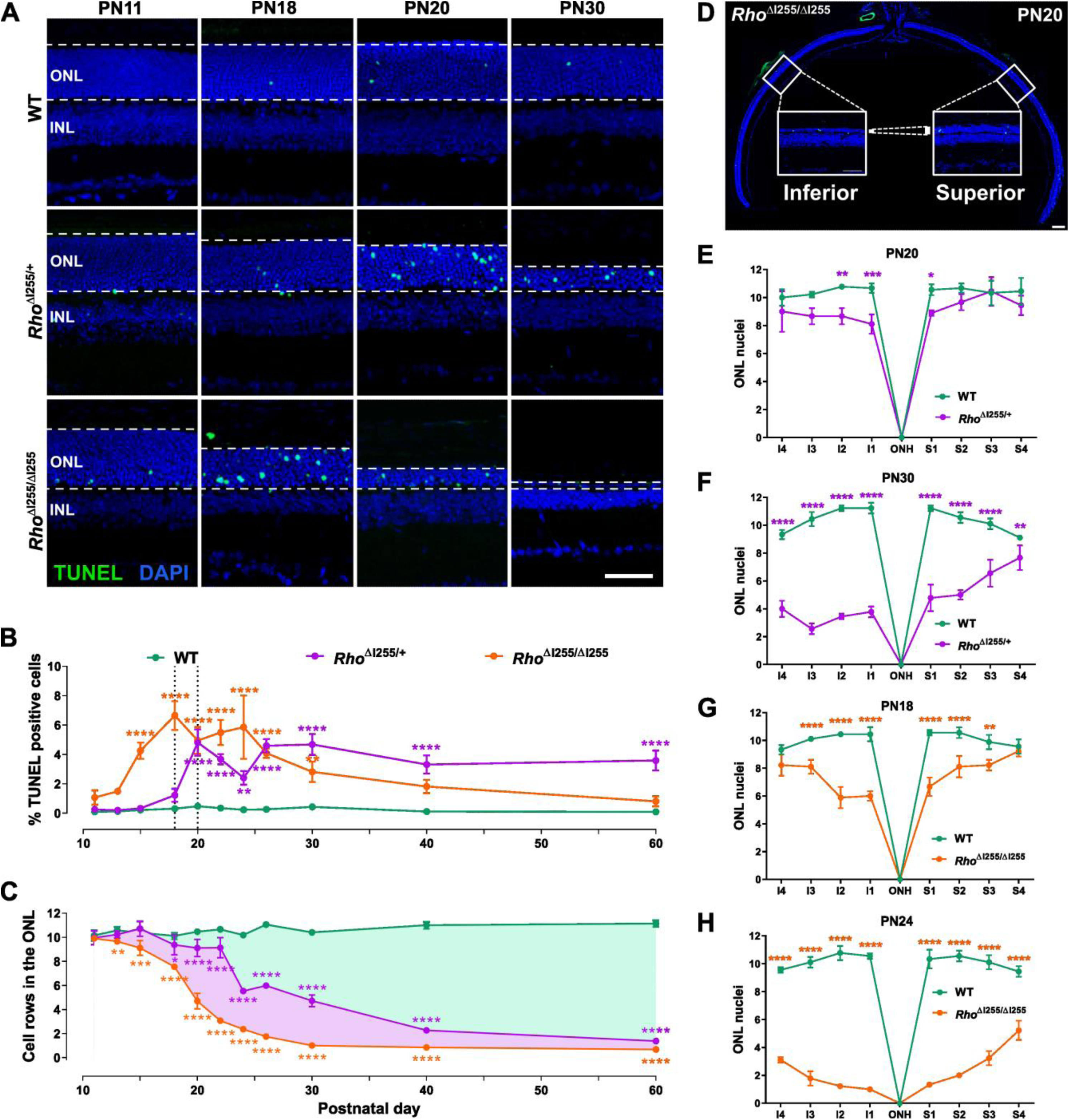
Temporal progression of photoreceptor cell death in *Rho*^ΔI255^ mice. (**A**) TUNEL staining (green) for dying cells in wild-type (WT), *Rho*^ΔI255/+^, and *Rho*^ΔI255/ΔI255^ retinae at postnatal (PN) days 11, 18, 20, and 30. Nuclear counterstain with DAPI (blue). (**B**) Percentage of cell death in outer nuclear layer (ONL) and (**C**) photoreceptor row counts between PN11 and 60 (n≥3). (**D**) Overview of retinal cross-section from PN20 *Rho*^ΔI255/ΔI255^ mice, superior-inferior direction, across optic nerve head. (**E**-**H**) Spider plots for ONL nuclear rows along the superior-inferior axis (n=3). (**E**) *Rho*^ΔI255/+^ retina at PN20 and (**F**) PN30. (**G**) *Rho*^ΔI255/ΔI255^ retinae at PN18 and (**H**) PN24 illustrate the regional variation in retinal degeneration (n=3). Scale bars: 50 µm (A and D), 100 µm (B); data expressed as mean ± SD; significance testing: Two-way ANOVA and Bonferroni’s multiple comparisons test; \**p* < 0.05, \*\**p* < 0.01, \*\*\**p* < 0.001, \*\*\*\**p* < 0.0001 (purple asterisks indicate differences between *Rho*^ΔI255/+^ and WT, orange asterisks represent significances between *Rho*^ΔI255/ΔI255^ and WT).

Conversely, the quantification of the rows of photoreceptor nuclei in the ONL revealed a drastic reduction of nuclei in homozygous and heterozygous mutants, with only approximately 4 and 9 remaining rows, respectively, at PN20. By PN30, in *Rho*^ΔI255/ΔI255^ mice, and at PN60, in *Rho*^ΔI255/+^ mice, only one row of nuclei was preserved in the ONL, whereas WT mice maintained approximately 10 to 11 rows of nuclei throughout the study period (Fig. 1C).

The observed photoreceptor loss in *Rho*^ΔI255^ mice showed a superior-inferior (dorsal-ventral) asymmetry, with more pronounced ONL row loss in the inferior retina than the superior retina. In the retina of PN20 *Rho*^ΔI255/+^, between 8.1 and 9.0 rows of nuclei were observed in the inferior retina, while between 8.8 and 10.4 ONL rows were present in the superior retina (Fig. 1E). At PN30, the number of photoreceptor cell rows ranged from 2.5 to 4.0 in the inferior region, and in the superior region, a range of 4.7 to 7.6 rows was maintained (Fig. 1F). In the homozygous *Rho*^ΔI255/ΔI255^ group, the average number of remaining nuclear rows in the inferior hemiretina ranged from 5.8 to 8.2 at PN18, while those in the superior hemiretina ranged from 6.6 to 9.2 (Fig. 1G). At PN24, this asymmetry was further increased, with only 1.0 to 3.1 rows remaining in the inferior area in contrast to 1.3 to 5.2 in the superior area (Fig. 1H).

This initial examination of the novel *Rho*_ΔI255_ model demonstrated a marked degenerative phenotype in heterozygous animals, in line with the dominant nature of human adRP caused by *Rho*_ΔI255_, and an even more expressed phenotype in homozygous animals.

### 2. *Rho*^ΔI255^ mice display RHO mislocalization

RHO is a major structural component of the outer segment (OS) discs and is critical for rod photoreceptor function [22–24]. Alterations in RHO production or distribution typically induce cytotoxic effects, leading to photoreceptor cell death [25–27]. To analyze underlying pathomechanisms further, we investigated RHO protein expression and distribution using immunofluorescence staining.

As early as PN11, mislocalization of RHO was observed in the ONL of heterozygous Rho^ΔI255/+^ and homozygous *Rho*^ΔI255/ΔI255^ mutant retinae. By PN15, a significant reduction in the length of photoreceptor outer segments (OSs) was observed in *Rho*^ΔI255/ΔI255^ mice (Fig. 2A, Fig. S1B). To quantify RHO distribution changes, the ratio of RHO mean fluorescence intensity (MFI) in the ONL and inner segment (IS) to that in OS was calculated using the following formula:

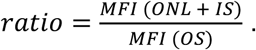

**Figure 2.**
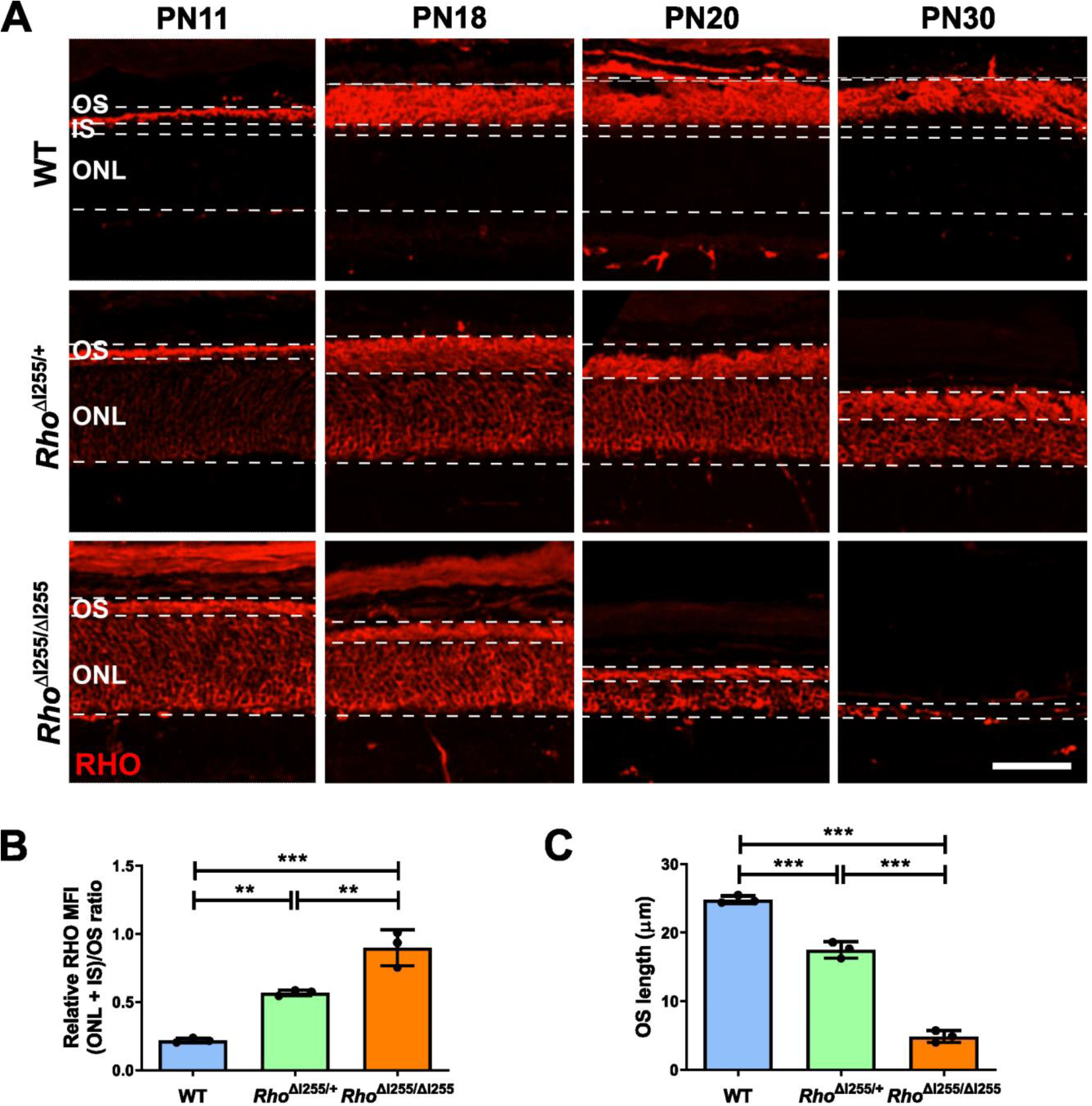
RHO mislocalization and reduced outer segment length in *Rho*^ΔI255^ retinae. (**A**) RHO immunostaining (red) in WT, *Rho*^ΔI255/+^, and *Rho*^ΔI255/ΔI255^ retina at PN11, 18, 20, and 30. (**B**) Ratio of RHO mean fluorescence intensity (MFI) between outer nuclear layer (ONL) and inner segment (IS) *vs.* outer segment (OS) ((ONL + IS)/OS)) at PN20 (n = 3). (**C**) OS length in WT, *Rho*^ΔI255/+,^ and *Rho*^ΔI255/ΔI255^ mice at PN20 (n = 3). Scale bars: 50 µm; data expressed as mean ± SD; significance calculated by two-way ANOVA and Bonferroni’s multiple comparisons test; \**p* < 0.05, \*\**p* < 0.01, \*\*\**p* < 0.001, \*\*\*\**p* < 0.0001.

At PN20, the ratio was significantly increased in both the *Rho*^ΔI255/ΔI255^ and the *Rho*^ΔI255/+^ mice, compared to age-matched WT mice, indicating a pronounced RHO mislocalization (Fig. 2B). Furthermore, analysis of OS length confirmed that *Rho*^ΔI255/ΔI255^ and *Rho*^ΔI255/+^ mice had significantly shorter OSs than WT mice, further highlighting the extent of photoreceptor degeneration (Fig. 3C).

**Figure 3.**
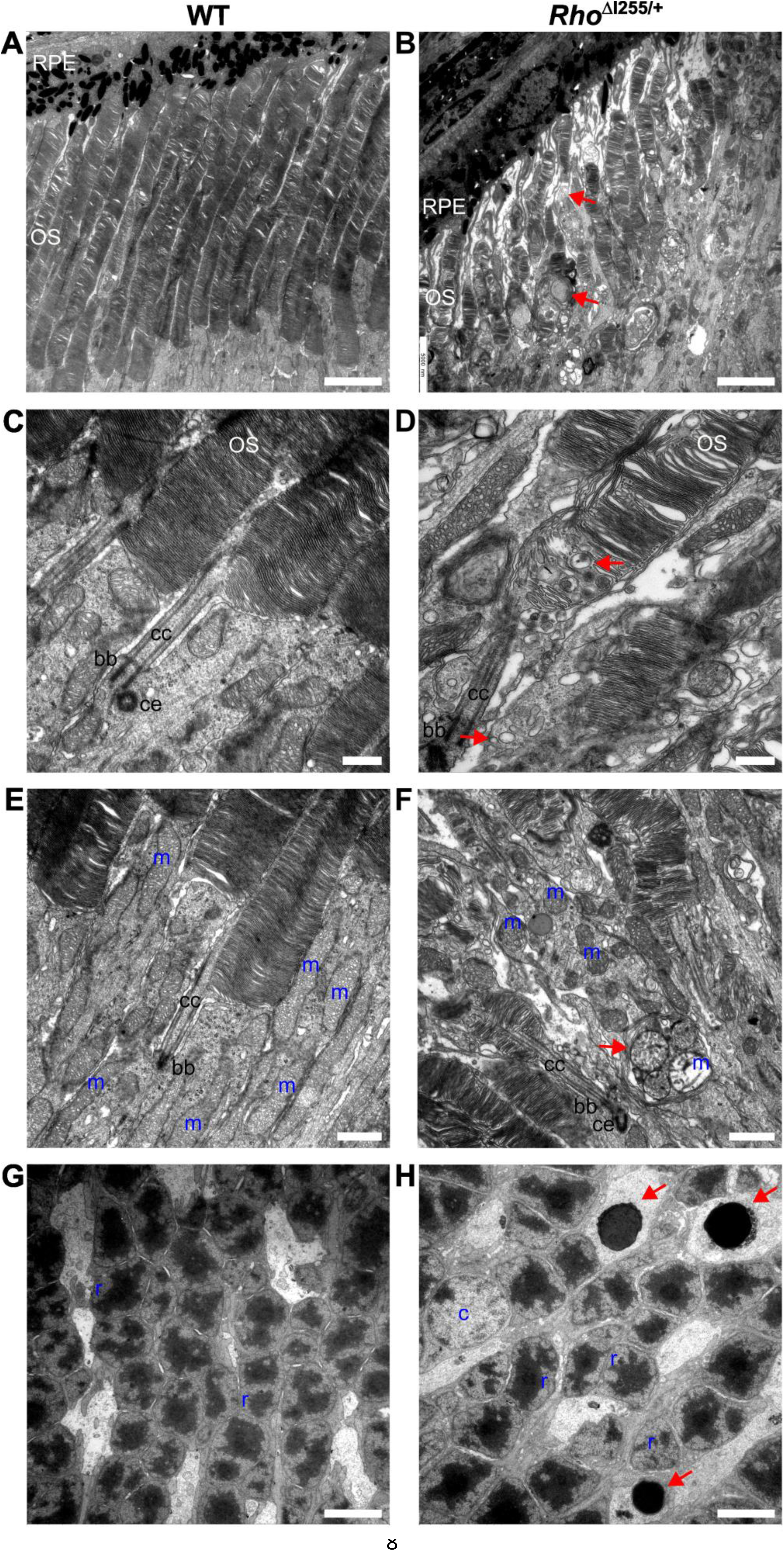
Transmission electron microscopy shows aberrant *Rho*^ΔI255/+^ rod photoreceptor segment formation. (**A-D**) Ultrastructure of retinal photoreceptors in WT and *Rho*^ΔI255/+^ mice. *Rho*^ΔI255/+^ photoreceptors displayed loose and irregularly shaped discs and vacuoles (red arrows). (**E**, **F**) Aberrant IS structures with smaller, round, swollen mitochondria (m) in the mutant retina compared to elongated mitochondria in the WT retina. Mitochondria were surrounded and enclosed by multiple membranes of autophagosomes, indicated by red arrows. (**G**, **H**) Close-up of outer nuclear layer (ONL). Pyknotic nuclei in *Rho*^ΔI255/+^ ONL (H, red arrows). Scale bars: 5000 nm (A, B, G, H), 500 nm (C, D), 1000 nm (E, F); RPE: retinal pigment epithelium, cc: connecting cilium, bb: basal body, ce: centriole, r: rod nuclei, c: cone nuclei.

These structural changes were analyzed in more detail by immunogold labeling and electron microscopy to determine the subcellular localization of RHO in the PN20 retina. In WT retinae, RHO was densely distributed across the well-structured rod OS (Fig. S2A, black arrows). A smaller amount of RHO was also detected within the rod inner segment (IS) of WT mice, which is consistent with expected normal protein synthesis patterns (Fig. S2C, black arrows). In contrast, a significant accumulation of immunogold-labelled structures was observed on the IS plasma membrane in *Rho*^ΔI255/+^ retinae (Fig. S2D, black arrows), which corroborated the findings of RHO mislocalization observed in immunofluorescence.

### 3. Photoreceptor segments are malformed and disorganized in *Rho*^ΔI255/+^ retinae

The impact of the *Rho*^ΔI255^ mutation on the photoreceptor OS was primarily observed through their shortening and the mislocalization of RHO in the ONL and IS, as revealed by RHO immunostaining. To investigate photoreceptor abnormalities induced by this mutation in greater detail, we examined the fine structure of photoreceptors in *Rho*^ΔI255/+^ retinae at PN20 by transmission electron microscopy (TEM). In healthy WT photoreceptors, OSs were elongated, intact, and correctly oriented in parallel bundles (Fig. 3A). In *Rho*^ΔI255/+^ mice, OSs were shortened, lacked correct orientation, and displayed variable-sized whorls (Fig. 3B, red arrows). At the ultrastructural level, WT mice had densely stacked, horizontally assembled discs in the OSs (Fig. 3C), whereas *Rho*^ΔI255/+^ mice exhibited disorganized, loose disc architecture, as well as multi-vesicular body-like structures (Fig. 3D, red arrows).

Given the crucial role of mitochondria in providing energy to maintain photoreceptor function, the ultrastructural changes in the IS were also assessed. The mitochondria within the IS of WT photoreceptors displayed an elongated morphology (Fig. 3E). However, in the IS of *Rho*^ΔI255/+^ photoreceptors, numerous damaged mitochondria with small, round, swollen shapes were observed, indicating mitochondrial stress and potential cell death (Fig. 3F). Many mitochondria were surrounded and enclosed by multiple membranes of autophagosomes, indicating mitophagy (Fig. 3F, red arrow).

In the ONL of WT mice, rod nuclei exhibited a round shape with a single clump of dark heterochromatin surrounded by lighter euchromatin. Meanwhile, in the ONL of *Rho*^ΔI255/+^ mutant retinae, some rod nuclei exhibited a round shape with a deep and dense structure (Fig. 3H, red arrows), characteristic of pyknosis and cell death [28].

### 4. Cone degeneration in *Rho*^ΔI255^ retina follows a characteristic spatiotemporal pattern

A defining feature of RP-type diseases is that the loss of rod photoreceptors eventually leads to the loss of cones (rod-cone dystrophy), accounting for the most significant part of visual impairment. To investigate pathological changes that affect cone photoreceptors in *Rho*^ΔI255^ mice, retinal cryosections were stained at different developmental stages with a cone arrestin antibody, labeling entire cone photoreceptors [29].

Signs of altered cone morphology were first observed at PN15 in *Rho^ΔI255/ΔI255^*mice and at PN20 in *Rho^ΔI255/+^* mice. Cone length was reduced over time, accompanied by signs of OS regression (Fig. 4A, Fig S3A). Further, a quantification of cone numbers per 100 µm along the vertical meridian of the retina revealed a progressive reduction in cone density (Fig. S3B). The morphological cone alterations in heterozygous *Rho*^ΔI255/+^ mice were observed by PN20, preceding the cone loss seen at PN30. This sequence of events is consistent with prior research on *rd1* mice, a widely used animal model of retinal degeneration [30]. The topographic assessment of cone degeneration in *Rho*^ΔI255/+^ retinae revealed a marked hemispheric asymmetry at both PN20 and PN30, with a significant reduction in cone numbers in the inferior retina (Fig. 4B-C).

**Figure 4.**
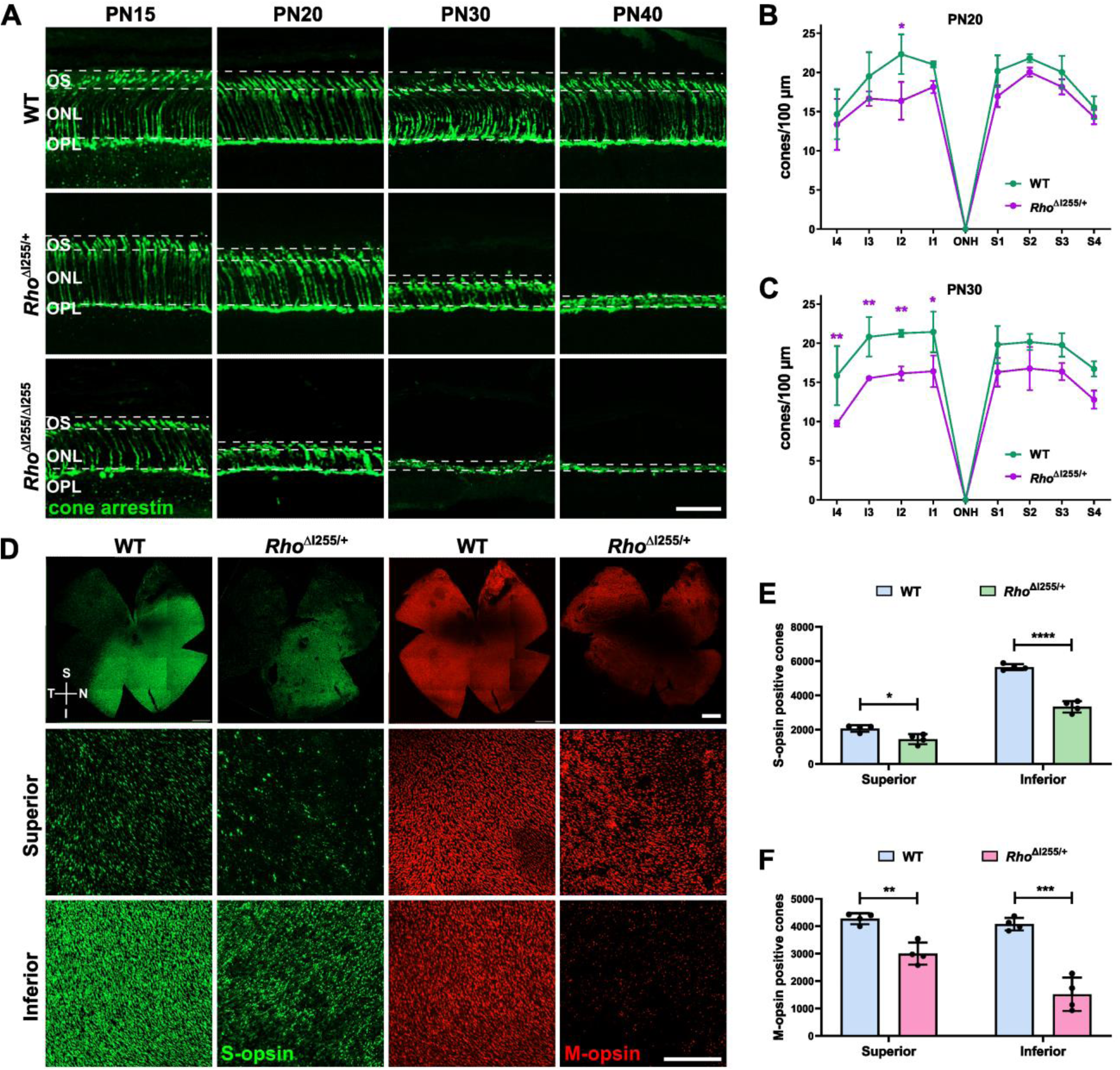
The *Rho*^ΔI255^ mutation leads to asymmetric cone degeneration. (**A**) Retinal cross-sections derived from PN15 to PN40 in WT, *Rho*^ΔI255/+^, and *Rho*^ΔI255/ΔI255^ animals. Immunolabeling for cone arrestin (green), showing progressive loss of cones. (**B**, **C**) Spider plots for cone density along the superior-inferior axis in WT and *Rho*^ΔI255/+^ mice at PN20 and PN30 (n = 3). (**D**) Wholemount immunostaining at PN30 in WT and *Rho*^ΔI255/+^ retina. S- and M-opsin immunolabeling suggested regional variations in cone cell loss. (**E**, **F**) Quantitative analysis of S- and M-opsin positive cones in the superior and inferior retina. PN30 counts of S- and M-opsin positive cells from analogous regions (30,000 µm^2^) of WT and *Rho^ΔI255/+^* retinae. Scale bars: 50 µm (A), 500 µM (D, 1^st^ row), 200 µM (D, 2^nd^ and 3^rd^ row); data expressed as mean ± SEM; significance calculated by two-way ANOVA; Bonferroni’s multiple comparisons test in B, C; unpaired two-tailed *t*-test in E, F; \**p* < 0.05, \*\**p* < 0.01, \*\*\**p* < 0.001, \*\*\*\**p* < 0.0001.

To more precisely quantify regional variations of cone loss, the retinal distribution of short- (S) and medium- (M) wavelength cone opsin was analyzed at PN30 via wholemount immunolabeling with anti-S-opsin and anti-M-opsin antibodies. As previously described for WT retina [31, 32], the M-opsin positive cone density followed a mild gradient, with higher numbers in the superior (dorsal) retina and lower numbers in the inferior (ventral) retina, while the S-opsin positive cone density followed a more distinct gradient between the inferior (high) and the superior (low) retina (Fig. 4D). For both opsins, a similar distribution pattern was observed in the *Rho*^ΔI255/+^ retina, although at lower densities. The analysis of S- and M-opsin densities in the superior and inferior retinal regions revealed marked disparity in the spatial degeneration profile (Fig. 4E-F). Specifically, S-opsin labelling in *Rho*^ΔI255/+^ retinae was significantly reduced in the superior retina (−30%) and inferior retina (−41%) (Fig. 4E). Similarly, M-opsin positive cones displayed a significant decrease in the superior retina (−30%) relative to the inferior retina (−63%) (Fig. 4F). Remarkably, S-opsin staining showed an ectopic redistribution from the OS to the cell bodies (Fig. S4B, white arrows). Furthermore, we noted an abnormal and swollen morphology with S-opsin staining in the inferior section of retinal wholemounts (Fig. S4B, white arrows), indicating that the OS had deteriorated and that therefore S-opsin could only be localized to the cell bodies. In the mutant retina M-opsin showed a similar distribution (Fig. S4F and H, white arrows). In conclusion, these findings indicate a geographical asymmetry of cone degeneration in *Rho*^ΔI255^ mice, with a pronounced effect observed for both S- and M-opsin positive cones in the inferior retina, a phenomenon which has also been observed for other RP mutations [33, 34].

### 5. Morphological changes in *Rho*^ΔI255^ outer and inner retinal neurons

Previous studies had shown that rod and cone photoreceptor cell death will eventually evoke atrophy and connective re-patterning of 2^nd^- and 3^rd^-order neurons in the retinal circuitry [35, 36]. To investigate a possible neuronal pathology in *Rho*^ΔI255^ mice, we examined morphological modifications in rod bipolar cells, horizontal cells (2^nd^-order neurons), amacrine cells (interneurons), and ganglion cells (3^rd^-order neurons) in *Rho*^ΔI255/+^ and *Rho*^ΔI255/ΔI255^ retinae at PN20, *i.e.,* close to or at the peak of photoreceptor loss.

Rod bipolar cells were identified using an antibody directed against the alpha isoform of protein kinase C (PKC-α) [37]. The cells displayed extensive ‘candelabra-like’ dendrites within the outer plexiform layer (OPL) (Fig. 5A, arrows) and distinct axonal end bulbs with terminal varicosities in the inner plexiform layer (IPL) of WT retinae (Fig. 5A, arrowheads). However, in *Rho*^ΔI255/+^ and *Rho*^ΔI255/ΔI255^ retinae, there were fewer dendrites within the OPL (Fig. 5A, white arrows) and terminal varicosities in the IPL appeared reduced in size (Fig. 5A, arrowheads), indicating degeneration of rod bipolar cell dendrites and axon terminals.

**Figure 5.**
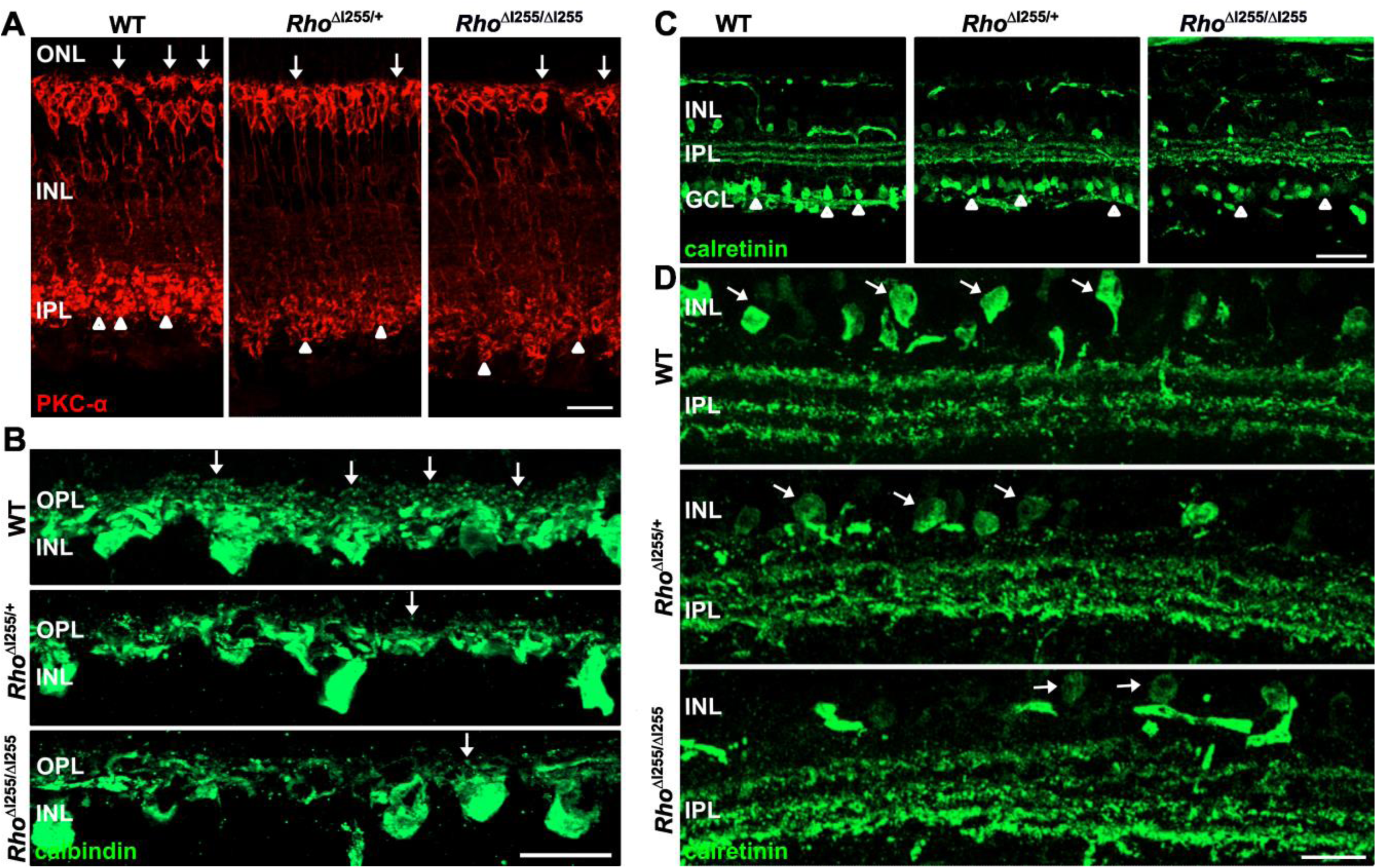
Inner retinal neurons in *Rho*^ΔI255^ mice display morphological alterations at PN20. Representative retinal sections from WT, *Rho*^ΔI255/+^, and *Rho*^ΔI255/ΔI255^ mice immunolabeled against PKC-α (red) (A) and calbindin (green) (B). Synaptic contacts with photoreceptors in rod bipolar cells (A, white arrows), horizontal cells (B, white arrows), and dendrites in rod bipolar cells (A, white arrowheads) were reduced in *Rho*^ΔI255/+^ and *Rho*^ΔI255/ΔI255^ mice. (C) Representative images of amacrine cells immunolabeled with anti-calretinin antibody (green) from WT, *Rho*^ΔI255/+^, and *Rho*^ΔI255/ΔI255^ retinal sections. (D) Higher magnification of calretinin staining indicated distortions in synaptic strata of the inner plexiform layer (IPL) in *Rho*^ΔI255/+^ and *Rho*^ΔI255/ΔI255^ retinae. Calretinin-positive cell numbers appeared lower (C, white arrowheads) and labeling weaker (D, white arrows) in the mutant inner nuclear layer (INL) and ganglion cell layer (GCL). Scale bars: 50 µm (C), 20 µm (A, B, D).

Horizontal cells, immunolabeled with an anti-calbindin antibody [37], exhibited a distinct punctate pattern with dendritic tips aligning at the superior border of the inner nuclear layer (INL) in WT mice. In contrast, in both *Rho*^ΔI255/+^ and *Rho*^ΔI255/ΔI255^ retinae, a decreased density of horizontal cell dendrites was observed, indicating a reduced connectivity (Fig. 5B, white arrows).

Amacrine cells, labeled with calretinin antibody [38], typically show terminals forming three distinct layers within the IPL and numerous cell bodies in the INL (Fig. 5D, white arrows) and ganglion cell layer (GCL; Fig. 5C, white arrowheads) of WT retinae [39]. However, the three strata in the IPL of *Rho*^ΔI255/+^ and *Rho*^ΔI255/ΔI255^ retinae were rather disorganized (Fig. 5D, second row), with both a reduction in stained amacrine cell bodies in the GCL (Fig. 5C, white arrows), and reduced calretinin immunolabeling in the INL (Fig. 5D, white arrows).

Taken together, these findings indicate a substantial structural disorganization of the neuroretina and a potential loss of synaptic function. The presence of morphological changes in 2^nd^- and 3^rd^-order neurons at PN20 in *Rho*^ΔI255/+^ mice (and even more pronounced in *Rho*^ΔI255/ΔI255^ mice) suggests a significant impact of the *Rho*^Δ*I255*^ mutation on retinal neuronal structure and connectivity beyond the outer retina also in respective adRP patients, which we suggest to verify in future work.

### 6. Increased Müller cell gliosis and microglial infiltration in the outer retina of *Rho*^ΔI255^ mice

Macro- and microglial cells play essential roles in maintaining retinal homeostasis, particularly in response to retinal diseases [40]. Astrocyte activation, reactive Müller cell gliosis, and microglial infiltration are common manifestations in retinal dystrophies [41–44]. We investigated the progression of these glial changes during retinal degeneration in *Rho*^ΔI255^ mutants, using immunostaining for glial fibrillary acidic protein (GFAP) and ionized calcium-binding adapter molecule 1 (Iba1). In WT retinae, the GFAP fluorescent signal was mainly restricted to the GCL (Fig. 6A). In contrast, in *Rho*^ΔI255/+^ (starting PN20) and *Rho*^ΔI255/ΔI255^ mice (from PN18), GFAP fluorescence extended to the IPL, INL, ONL, and OPL (Fig. 6A). Analysis of MFI from GCL to subretinal space confirmed that GFAP was significantly increased in mutants compared to WT mice (Fig. 6B, C).

**Figure 6.**
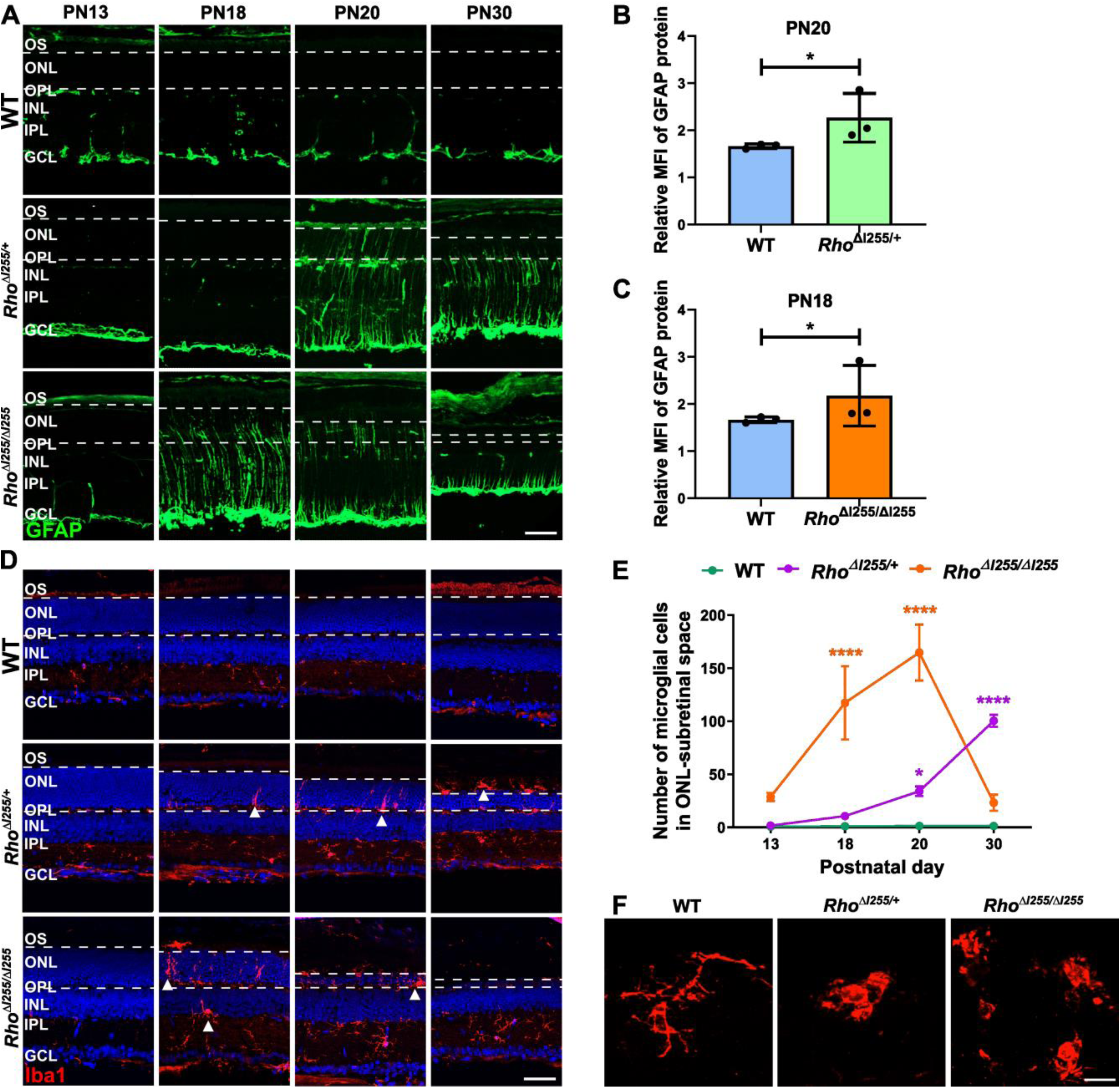
Müller cell and microglia activation in mutant *Rho*^ΔI255^ mice. (**A**) Immunolabeling for GFAP (green) in WT, *Rho*^ΔI255/+^, and *Rho*^ΔI255/ΔI255^ retina between PN13 and PN30. (**B**, **C**) MFI analysis of GFAP protein in retinae of *Rho*^ΔI255/+^ mice at PN20 and *Rho*^ΔI255/ΔI255^ mice at PN18, compared to WT (n = 3). (**D**) Iba1 staining (red) in WT and mutant retina from PN13 to PN30, counterstained with DAPI (blue). White arrows represent microglia within the INL; white arrowheads indicate ONL and subretinal infiltration. (**E**) Quantification of microglial cell bodies within the ONL and subretinal space (n = 3). (**F**) Iba1 positive microglia morphology in representative high-magnification images of WT (PN20), *Rho*^ΔI255/+^ (PN30), and *Rho*^ΔI255/ΔI255^ (PN20) mutant retina. Scale bars: 50 µm (A, D), 10 µm (F); data expressed as mean ± SEM; significance calculated by two-way ANOVA and Bonferroni’s multiple comparisons test. \**p* < 0.05, \*\*\*\**p* < 0.0001.

While in WT mice, Iba1-positive cells were primarily observed in the inner retinal layers, particularly the GCL and IPL (Fig. 6D), Iba1 staining in *Rho*^ΔI255^ mice showed microglial infiltration in the outer retina. In *Rho*^ΔI255/+^ and *Rho*^ΔI255/ΔI255^ retinae, Iba1-positive cells migrated to the INL, ONL, and subretinal space (Fig. 6D, white arrowheads). Quantitative assessment of Iba1-positive cells in ONL and subretinal space revealed a significant increase in microglial infiltration in *Rho*^ΔI255/+^ retinae at PN20 and *Rho*^ΔI255/ΔI255^ retinae at PN18 (Fig. 6E). Microglia morphology in WT retinae indicated a predominantly resting state, with small cell soma, minimal perinuclear cytoplasm, and numerous fine, highly branched processes (Fig. 6F). In contrast, in *Rho*^ΔI255/+^ and *Rho*^ΔI255/ΔI255^ mice numerous amoeboid microglial cells were observed in different layers of the retina. These cells were identified by their enlarged somas and thick retractive processes, indicating activated microglia (Fig. 6F) [45].

These results show that Müller cell gliosis and microglial activation in the outer retina are prominent characteristics of *Rho*^ΔI255^ mutation-induced retinal degeneration as is typical for RP.

### 7. *In vivo* retinal morphology and function are severely compromised in *Rho*^ΔI255/+^ mice

The *ex vivo* examinations thus far indicated that the *Rho*^ΔI255^ mutation induces a severe retinal degenerative phenotype. To assess possible macromorphological alterations in *Rho*^ΔI255/+^ mice, we then used *in vivo* imaging. At one month of age, native images of the fundus of *Rho*^ΔI255/+^ mice displayed a leopard skin-like appearance (Fig. 7A), indicative of severe photoreceptor degeneration and a loss of the regular photoreceptor/RPE interface. Optic coherence tomography (OCT) revealed a corresponding massive reduction in retinal thickness, which at PN30 was essentially limited to the outer retina (Fig. 7B, Fig. S4).

**Figure. 7.**
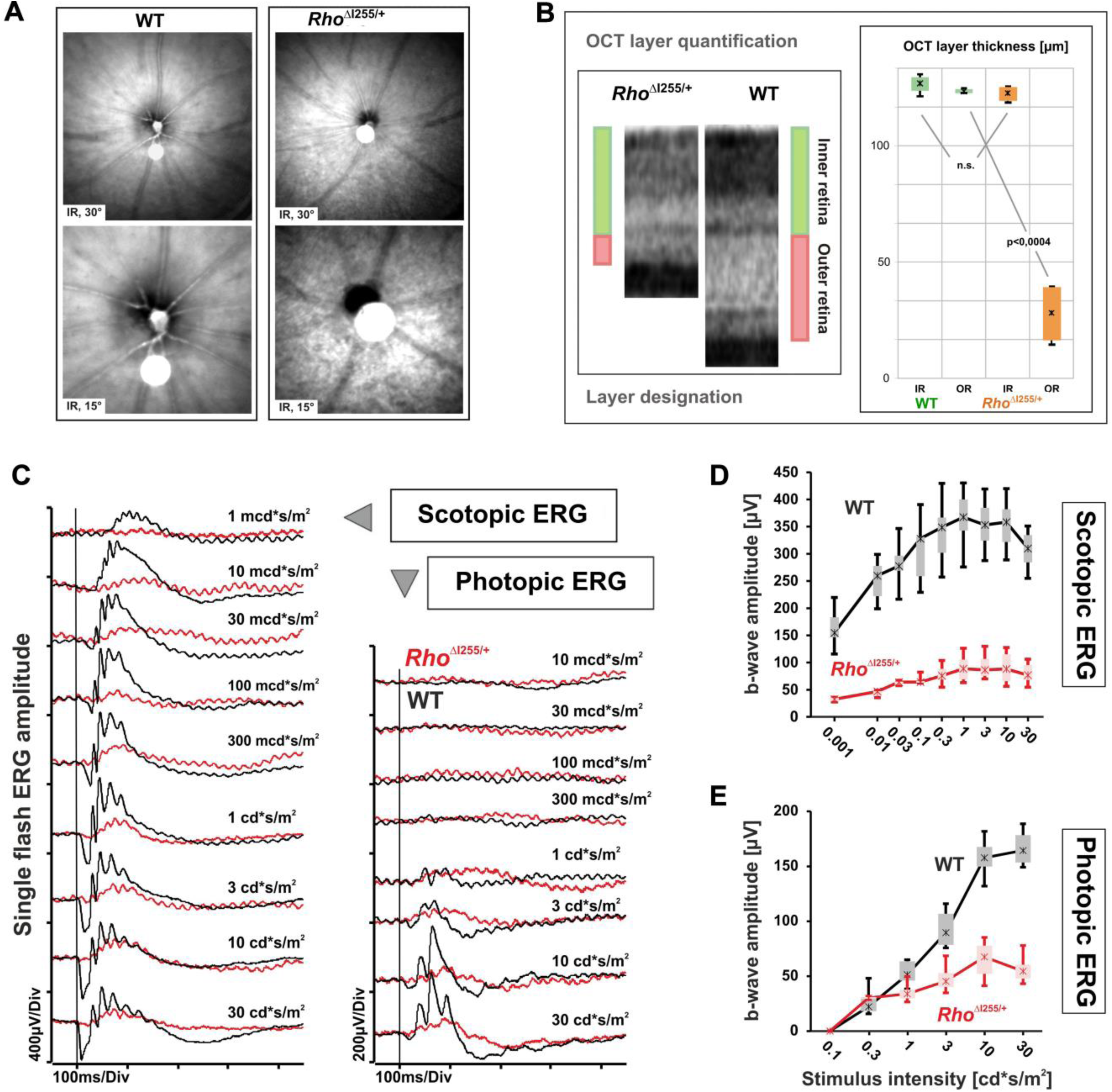
*In vivo* retinal morphology and function of the *Rho*^ΔI255/+^ line at PN30. (**A**) Representative native infrared (IR) scanning laser ophthalmoscopy (SLO) fundus images at two magnifications (device setting of 30° or 15°). (**B**) Representative OCT sections and quantification of layer contribution to the loss of retinal thickness in OCT. Data on the right are shown as box-and-whisker plots for WT and *Rho*^ΔI255/+^ mice (n = 4 eyes per group). Boxes: 25–75% quantile range, whiskers: 5% and 95% quantiles, asterisks: median. A one-sided t-test for two samples with unequal variances was used to calculate the *p*-value. (**C**) Representative dark-adapted (scotopic) single flash intensity series ranging from 1 mcd*s/m^2^ to 30 cd*s/m^2^ (left image) and light-adapted (photopic) single flash intensity series ranging from 10 mcd*s/m^2^ to 30 cd*s/m^2^ on a 30 cd/m^2^ background (right image); black: WT, red: *Rho*^ΔI255/+^. The vertical lines indicate the time of the light stimulus. (**D, E**) Quantitative evaluation of scotopic (D) and photopic (E) ERG b-wave amplitudes (box-and-whisker-plots) for WT and *Rho*^ΔI255/+^ mice (n = 10 eyes per group). Boxes: 25–75% quantile range, whiskers: 5% and 95% quantiles, asterisks: median.

As a next step, we assessed retinal function with electroretinography (ERG) to analyze the impact of morphological defects on retinal functionality. At PN30, a dark-adapted ERG intensity series revealed extremely low scotopic responses of *Rho*^ΔI255/+^ mice compared to WT controls (Fig. 7C, D). In particular, the a-waves that are typically dominated by rod outer segments were barely visible in *Rho*^ΔI255/+^ mutants throughout the ERG intensity series. In conjunction with the shape of the waveform and the low amplitude at 10 mcd*s/m^2^ (below which ERG responses are purely rod-driven), this indicated that the scotopic ERG in *Rho*^ΔI255/+^ mice at this age was mainly cone-driven, as often seen in RP. Further, the photopic ERG of *Rho*^ΔI255/+^ mice were found to be reduced (Fig. 7C, E), implicating that cone system responses were already affected at this age. Our result indicates a much earlier secondary cone degeneration in the *Rho*^ΔI255/+^ mouse than in the *Rho*-knockout mouse [46, 47], suggesting that the presence of a pathological rhodopsin variant is more deleterious than a complete absence of the protein, although both may lead to a lack of rod outer segments.

### 8. *Rho*^Δ*I255*^ mutant retina undergoes apoptotic and non-apoptotic cell death

To identify potential targets for future therapy development in *Rho*^ΔI255^-induced photoreceptor loss, a thorough understanding of the underlying neurodegenerative mechanisms is essential. In our hands, a key to the degeneration in the *Rho*^ΔI255^ mouse model is the type of cell death driving photoreceptor loss. Consequently, we analyzed several different markers for processes typically associated with either apoptosis or non-apoptotic cell death [19]. In this context it is worth to note that TUNEL assays label a large variety of different cell death modalities and do not *per se* allow direct inferences on underlying degenerative mechanisms [48].

To test specifically for apoptotic cell death, we first stained for cleaved, activated caspase-3 in WT, *Rho*^ΔI255/+^, and *Rho*^ΔI255/ΔI255^ retinae [49] at different time points ranging from PN11 to PN60 (Fig. 8A, B). While the WT retina exhibited minimal caspase-3 activity, the *Rho*^ΔI255/ΔI255^ retina displayed a peak of such cells at PN18. In the heterozygous mutant, the peak of caspase-3 activity was delayed to approximately PN40 (Fig. 8B). However, especially at the onset of degeneration in the homozygous and heterozygous mutants, there was a mismatch between the numbers of caspase-3 positive cells and the amounts of dying cells detected with the TUNEL assay (*cf*. Fig. 1), suggesting that caspase-independent, non-apoptotic cell death might play an important role in *Rho*^ΔI255^ photoreceptor loss.

**Figure 8.**
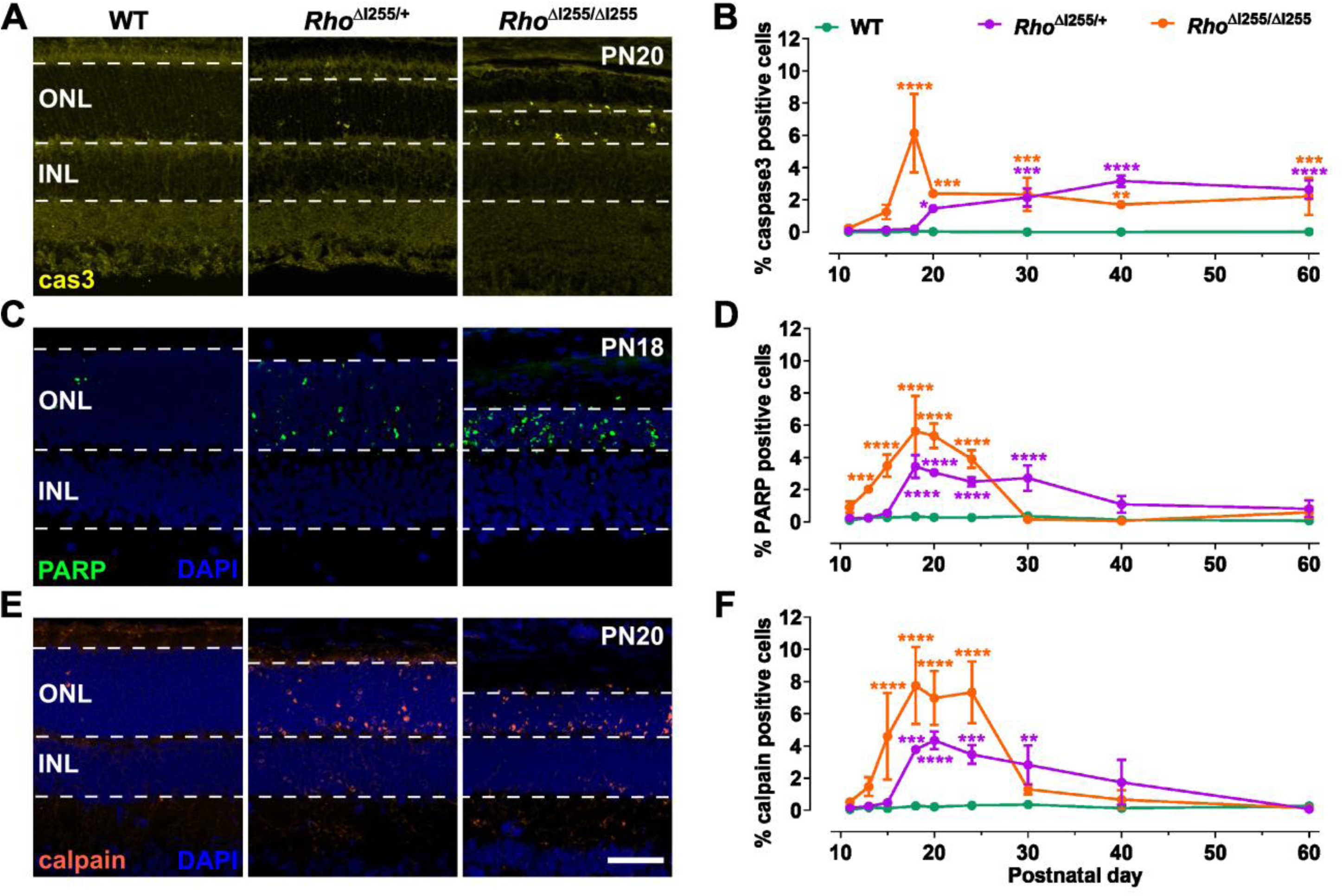
Apoptosis and non-apoptotic cell death in *Rho*^ΔI255^ retina. Immunostaining and activity assays for different cell death markers were performed on retinal cross-sections derived from WT, as well as *Rho*^ΔI255/+^ and *Rho*^ΔI255/ΔI255^ mutants. (**A**) At PN20, immunostaining for cleaved, activated caspase-3 (cas3; yellow) was used to detect classical apoptosis in WT and mutant genotypes. (**B**) Quantification of caspase-3 positive cells in the outer nuclear layer (ONL) of WT, *Rho*^ΔI255/+^, and *Rho*^ΔI255/ΔI255^ retina. (**C**) PARP activity (green) in WT and mutant retina at PN18. (**D**) Quantification of PARP activity-positive cells in WT and mutant ONL. (**E**) Calpain activity (orange) in WT and mutant retina at PN20. (**F**) Quantification of calpain activity-positive cells in WT and mutant ONL. Scale bar: 50 µm. Data expressed as mean ± SD. Significance calculated by two-way ANOVA and Bonferroni’s multiple comparisons test. \**p* < 0.05, \*\**p* < 0.01, \*\*\**p* < 0.001, \*\*\*\**p* < 0.0001.

To explore non-apoptotic photoreceptor cell death, we initially used an *in situ* activity assay for poly(AD-ribose)polymerase (PARP) performed on unfixed retinal tissue sections (Fig. 8C) [50]. While the WT retina did not show appreciable amounts of PARP activity-positive cells in the ONL at any time point studied, the homozygous *Rho*^ΔI255/ΔI255^ mutant, from PN13 onwards, showed an early and relatively strong increase in PARP activity-positive ONL cells. In heterozygous *Rho*^ΔI255/+^ retina, PARP activity began at a later stage, at PN18, and appeared to be sustained for a more extended period of time (Fig. 8D). Similar to PARP, the activity of Ca^2+^-dependent calpain-type proteases has previously been linked to non-apoptotic retinal degeneration [51], and we employed an *in situ* assay to resolve this activity in photoreceptors. From PN18 onwards, the number of calpain activity-positive cells was significantly increased in the retina of heterozygous *Rho*^ΔI255/+^ mice compared to WT. In contrast, calpain activity-positive cells appeared earlier and at a higher number in the retinae of homozygous *Rho*^ΔI255/ΔI255^, reaching a peak at PN18 (Fig. 8E, F).

Taken together, in both homozygous and heterozygous *Rho*^ΔI255^ retinae, an activation of caspase-3, a marker for apoptosis, as well as PARP and calpain, markers for non-apoptotic cell death, were observed and may serve as targets in future work on adRP. These elevated enzyme activities coincided with the period of the most extensive photoreceptor cell death (Fig. 1B), indicating that both apoptotic and non-apoptotic mechanisms were causally involved in *Rho*^ΔI255^ retinal degeneration. Relative to each other, calpain activity appeared earlier and was numerically more important than caspase-3 activation.

## Discussion

The causative mechanisms underlying human *RHO*^ΔI255^-related adRP remain largely unclear and their elucidation, as well as therapy developments, necessitate *in vivo* disease models that closely mimic the human condition. In the present work, we have generated and characterized a new knock-in mouse model homologous for the dominant human *RHO*^ΔI255^ RP mutation. The new *Rho*^ΔI255^ mouse faithfully reproduces a number of key pathological features of human adRP, including photoreceptor loss, inner retinal remodeling, and evidence for neuroinflammation, and may prove instrumental for future therapy developments.

### Retinal disease pathogenesis caused by *RHO* mutations

A large variety of mutations in the human *RHO* gene can cause adRP and according to retinal dystrophy patterns and clinical manifestations these are categorized into two main classes [2]: Class A is characterized by early-onset severe rod dystrophy, while class B presents with a later-onset and milder phenotype, progressing more slowly. Subclass B1 relates to patients suffering from the T17M, P23H, T58R, V87D, G106R, and D190G mutations, photoreceptor degeneration tends to exhibit an inferior to superior regional predisposition. This conforms to what is seen for the *RHO*^ΔI255^ mutation, which may thus also be categorized within subclass B1 [8, 11, 13].

In analogy to human adRP patients, *Rho*^ΔI255^ mice suffer from a progressive retinal degeneration leading to a severe disruption of retinal structures and eventually to an impairment of visual function up to blindness. In the novel adRP model, we found superior-inferior variations in retinal degeneration, with inferior retinae degenerating more rapidly than superior retinae. Similar superior-inferior (dorsal-ventral) differences in retinal structure have also been documented in other animal models of mutation-induced retinal degeneration [52–54]. Mutated RHO^ΔI255^ protein is found to accumulate in the perinuclear and IS regions of retinae in in both the homozygous *Rho*^ΔI255/ΔI255^ and heterozygous *Rho*^ΔI255/+^ mice, preceding photoreceptor cell death (*cf*. Fig. 2). Although mislocalization alone does not induce aggregation [55], previous *in vitro* studies have shown that *Rho*^ΔI255^ forms multiple, variably sized aggregates [17]. These findings suggest that mislocalized RHO^ΔI255^ aggregation not only precedes but also contributes to photoreceptor cell death, consistent with recent *in vivo* observations in P23H and G188R mutants [56]. Further, RHO aggregates were found to impair the ubiquitin-proteasome system [57, 58], and strategies aimed at reducing or eliminating RHO aggregates have been demonstrated to delay retinal degeneration in rodent models [59–62].

*Rho*^ΔI255^ retinae also exhibited decreased OS length, which may be attributed to two possible mechanisms: 1) *Rho*^ΔI255^ may induce misfolded RHO protein aggregation in the ONL and IS, preventing RHO transport to the OS. 2) Mislocalized RHO^ΔI255^ aggregates may exert a dominant-negative effect by trapping the mature form of normal RHO, preventing it from reaching the OS. In previous *in vitro* studies, we produced RHO^ΔI255^ as an EGFP fusion protein which sequestered RHO^WT^-mcherry into perinuclear aggregates. This prevented correct RHO localization in photoreceptors, suggesting a dominant-negative effect for the mutant protein [17]. The expression of RHO^ΔI255^ in rod photoreceptors *in vivo* also caused perinuclear aggregates and impaired outer segment formation. However, distinguishing normal from mutant RHO *in vivo* remains difficult as both forms differ only by a single amino acid.

Previous research has shown that rods in heterozygous *Rho*-knockout mice were characterized by slight changes in OS disc diameter and volume [63], but there was no degeneration comparable to that of RHO^ΔI255^ mutants. In contrast, homozygous *Rho*-knockout mice fail to develop rod OS [64]. However, even in those animals cone degeneration and associated loss of function sets occur as late as about postnatal weeks 7-8, followed by the loss of the final continuous layer of rod cells [65]. The fact that the secondary cone degeneration in RHO^ΔI255^ mice had started much earlier than in the *Rho*-knockout mouse, provides evidence that the presence of a pathological rhodopsin variant is more deleterious than a complete absence of any RHO protein and even a total lack of rod outer segments. Although the concept of an increasing expression of WT RHO has been found to show some promise in animal models of adRP [66] and arRP [67], we feel in the light of our data that rather a decreased expression of rhodopsin might alleviate the proteostatic stress on rod photoreceptors, which is also supported from recent work [27].

We have observed regional variations in cone loss, including S- and M-opsin positive cones, with the inferior retina degenerating more rapidly than the superior retina. This finding very well matches the “bystander effect” characteristic for RP and proposing that cones primarily unaffected by a disease die whenever rod cells are lost to a sufficient extent.

### The *Rho*^ΔI255^ mutation affects retinal function and inner retinal structure

Retinal function in *Rho*^ΔI255/+^ animals was consistent with a model of adRP. There was no clear sign of any rod functionality in the ERG at postnatal day 30 and beyond, including the lack of a-waves in the scotopic ERGs, while cone function declined subsequently to rod cell loss over time, manifested in reduced cone-driven responses both in the scotopic and photopic records. The same sequence may also be found in human RP patients, however obviously at a different time scale. Regarding the more distal visual pathway, we discovered that the incipient photoreceptor loss led to a significant anatomical remodeling of the downstream neuroretina. This affected 2^nd^- and 3^rd^-order neurons, including bipolar cells and the amacrine cells of the horizontal layer. Strikingly, this appeared concomitant to the process of photoreceptor degeneration. Waite and Crag had described as early as in 1979 [68] that the loss of primary sensory neurons caused by destroying the whiskers of rats resulted in a rapid loss of 2^nd^- and 3^rd^-order cerebral neurons as a consequence of the loss of sensory input. Auditory hair cell loss in the cochlea of mice was recently shown to cause altered primary auditory neuron functions, and latent neurite retraction at the hair cell-auditory neuron synapse [69]. In retinal disorders, mutant photoreceptors fail to deliver appropriate input to the inner retina [35]. However, the complete loss of photoreceptors after degeneration in RHO^ΔI255^ mice does initially not change thickness and layering of the inner retina, but rather disrupts structural and functional connectivity of the inner retinal circuitry instead. Similar results have been obtained in other RP rodent models, such as the *rd10* mouse [70], the rd1 (*rd*/*rd*) mouse [71], and the P23H rat [37]. As many details of inner retinal functional remodeling in human tissue still remain unclear, we believe that further investigations in the *Rho*^ΔI255^ lines will help to shed some light also on the downstream retinal alterations.

### Müller cell gliosis and markers of neuroinflammation in *Rho*^ΔI255^ retina

A major marker of retinal gliosis, GFAP, was upregulated in Müller cells of *Rho*^ΔI255^ mice upon degeneration, which may be associated with Müller cell hypertrophy [71]. Additionally, resident microglial cells in the inner retina were activated and migrated towards the outer retina. We found their highly arborescent morphology transformed to an amoeboid shape, accumulating in the ONL and subretinal space, potentially acting as phagocytes and clearing dying cells, as previously described [72].

Reactive gliosis, microglial activation and neuroinflammation are hallmarks of central nervous system neurodegenerative diseases [73]. However, whether these cellular responses are causative or consequential to neuronal damage and whether they are beneficial or detrimental to photoreceptor survival is still a matter of debate [36]. In our studies on *Rho*^ΔI255/+^ and *Rho*^ΔI255/ΔI255^ mice, microglia migrated into the INL and ONL at PN20 and PN18, respectively, *i.e.,* about 2 days after the onset of photoreceptor cell death. This temporal sequence underlines that microglial activation was not causing photoreceptor cell death and that, instead, microglia migrated into the ONL to clear away dying cells and debris. Other studies have reported that microglial phagocytosis promotes neuronal death by engulfing stressed but viable photoreceptors, a process that they termed phagoptosis [74, 75]. Chronic microglial activation can also release noxious factors that mediate neuroinflammation and pro-apoptotic events [35]. Moreover, activated microglia cells can stimulate Müller cells to release potentially neurotoxic cytokines such as interleukin (IL)-1β [76], synergistically promoting chronic neuroinflammation and retinal cell death. Müller cell activity, on the other hand, may also protect neurons by releasing neurotrophic factors, secreting antioxidants, and degrading excitotoxins [77]. It appears likely that microglial activation and Müller cell gliosis are a response to injury mechanism that primarily plays a protective role in the early stages of retinal degeneration but may become detrimental in chronic retinal dystrophy [78], which is a further course of events that is in current belief typical for the RP group. In addition, astrocytes and Müller cells contribute to promote retinal neovascularization by releasing vascular endothelial growth factor (VEGF) [79, 80]. However, at the fundus imaging level, we did not observe significant vascular atrophy or neogenesis in our animal model.

### Cell death mechanisms underlying *Rho*^ΔI255^-dependent photoreceptor loss

Apoptosis, the best-characterized form of programmed cell death, has long been thought to be the main mechanism in photoreceptor loss [81]. Yet, apoptosis may be more related to developmental processes and retinal maturation rather than mutation- or disease-induced degeneration [82], and research from the last decade indicated that photoreceptor cell death is more likely to be governed by non-apoptotic mechanisms, both in recessive RP and in adRP [19, 83].

In our TEM examination, *Rho*^ΔI255/+^ photoreceptors displayed characteristic features of pyknosis, *i.e*. round-shaped and noticeably denser nuclear structure indicative of chromatin condensation [84]. However, pyknosis cannot be considered a defining criterion, since it is seen in both apoptotic and non-apoptotic cell death [85], including in degenerating photoreceptors in RP [86]. In the inner segment, mitochondria were surrounded and enclosed by multiple membranes of autophagosomes, indicating mitochondrial stress and mitophagy. In the *Rho*^ΔI255^ retina this will likely reduce ATP production and contribute to retinal degeneration, as has already been demonstrated for different types of retinal diseases [87]. Importantly, a lack of ATP would prevent the execution of ATP-dependent apoptotic cell death, notably the ATP-dependent activation of so-called executioner caspases such as caspase-3 [85].

Nevertheless, a subset of *Rho*^ΔI255^ mutant photoreceptors exhibited activation of caspase-3, *i.e.* a cysteine-protease which plays a central role in executing apoptosis. While this finding would suggest the execution of apoptosis, the numbers of ONL cells exhibiting activated caspase-3 do not correspond to the numbers of cells positive for the TUNEL assay and appear overall too low to explain the rapid progression of photoreceptor loss.

Importantly, we found an early activation of calpain, *i.e.* a Ca^2+^-dependent cysteine-protease often connected to non-apoptotic cell death [88]. Calpain activity typically is associated with PARP activity and this is true also for photoreceptor death in other RP models, such as the P23H mutation in the *Rho* gene, different mutations in the *Pde6* gene (*rd1, rd10, V685M,* and *D670G)*, as well as the *Rpe65 KO* in the *RPE65* gene ([19, 20, 81]. PARP activity can consume exceedingly high levels of NAD^+^ leading to ATP depletion and a form of cell death termed PARthanatos [89]. Since the execution of caspase-dependent apoptosis requires ATP [90], PARthanatos and apoptosis seem to be axiomatically incompatible with each other. In other words, a cell that has activated PARP can no longer undergo apoptosis. In photoreceptors, PARthanatos shows a mechanistic overlap with cGMP-dependent cell death [83] in animal models for recessive RP, and future research may determine to what extent this is true also for retinal degeneration caused by the *Rho*^ΔI255^ mutation.

Taken together, the fact that we found the concomitant activation of caspase-3 and calpain/PARP, similar to what was seen earlier in the transgenic *Rho*^S334ter^ rat model for adRP [19, 91] suggests that at least two different cell death pathways can be triggered by the *Rho*^ΔI255^ mutation. Likely, these distinct pathways are executed separately in different subpopulations of photoreceptors. For an effective therapeutic development for adRP this may mean that different pathways need to be addressed simultaneously in a form of combination therapy [92], similarly to what is nowadays the norm for the treatment of cancer.

## Conclusion

The novel *Rho*^ΔI255/+^ *in vivo* model of autosomal dominant retinitis pigmentosa presented here displays a strong degenerative phenotype at several levels and provides thus evidence that the homologous human form of adRP may be seen as a systems disorder of the neuroretina affecting not only photoreceptors but the overall retina already early on in the course of degeneration. Further, several distinct degenerative mechanisms were identified to govern the primary photoreceptor degeneration, which in turn opens more than a single avenue for the future development of potential mutation-independent therapies.

## Materials and Methods

### Generation of the *Rho*^ΔI255^ mouse model and animal use

The healthy wild-type (WT) human and mouse RHO protein harbors two consecutive isoleucine residues at positions 255 and 256. One codon for one of these isoleucine (I) residues is missing in the human disease-causing allele [8]. The I255 residue is encoded in exon 6 of the *RHO* gene and located near the N-terminus of the sixth transmembrane domain of the RHO protein. To recreate the ΔI255 (c.768_770delCAT) mutation in the mouse, a knock-in strategy was used to insert a codon deletion equivalent to ΔI255. This mouse model was generated by the company GenOway (Lyon, France) using a standard homologous recombination technique in murine embryonic stem (ES) cells and C57BL/6J mice (JAX stock #000664) as genetic background. ES cell clones were genotyped by both PCR and Southern blot analysis, and the presence of the mutation was verified by sequencing. One fully characterized ES clone was used for injection into blastocysts of C57BL/6J mice to generate chimeric animals. Animals displaying germ-line transmission were subsequently bred to homozygosity. The experiments used either animals from the homozygous mouse line (*Rho*^ΔI255/ΔI255^) or heterozygous animals (*Rho*^ΔI255/+^) that were obtained by crossing homozygous mice with C57BL/6J wild-type (WT) animals.

Animals were housed in the Tübingen Institute for Ophthalmic Research animal facility under standard white cyclic lighting, had free access to food and water, and were used irrespective of gender. All procedures for animal studies were performed in accordance with the Association for Research in Vision and Ophthalmology (ARVO) declaration for the use of animals in ophthalmic and vision research and the law on animal protection issued by the German Federal Government (Tierschutzgesetz) and were approved by the institutional animal welfare office of the University of Tübingen under the registration No. AK02/20M. All efforts were made to minimize the number of animals used and their suffering.

### Histology and immunofluorescence staining

Animals from different postnatal days (PN) were sacrificed, and their eyes were enucleated and fixed in 4% paraformaldehyde (PFA) in 0.1 M phosphate buffer (pH 7.4) for 45 minutes (min) at 4 °C. PFA fixation was followed by cryoprotection in graded sucrose solutions (10, 20, 30%). Unfixed eyecups were directly embedded in cryomatrix (Tissue-Tek, Leica, Bensheim, Germany). Sagittal 14 µm sections were obtained and stored at −20 °C. Sections were incubated overnight at 4 °C with different primary antibodies (Table S1). Immunofluorescence was performed using Alexa Fluor 488 or 568 dye-conjugated goat anti-mouse or anti-rabbit antibodies (Table S1). For imaging, sections were mounted with Fluoromount-G Mounting Medium (EMS, 17984-25).

### Retinal wholemount staining

Following the previous protocol, retinae from WT and *Rho*^ΔI255/+^ mice at PN30 were collected for wholemount staining [93]. In brief, after corneal perforation, eyes were prefixed in 4% PFA for 5 min, then the anterior segment and vitreous bodies were removed, and eyeballs were fixed in 4% PFA for an additional 25 min. Fixed retinae were dissected from the pigmented epithelium and washed in bleaching solution (0.5% KOH and 3% H_2_O_2_ dissolved in phosphate-buffered saline; PBS) for 20 min, 6 times at room temperature (R.T.) to remove pigment. Retinae were washed with PBS 3 times and incubated in normal donkey serum (NDS) blocking solution at 4 °C overnight. Then, retinae were incubated for 3 days with S-opsin and M-opsin primary antibodies and for 2 further days with donkey anti-goat or anti-rabbit secondary antibodies (Table S1). Stained retinae were flat-mounted and covered with coverslips, with the photoreceptor layer facing upward.

### TUNEL assay

Terminal deoxynucleotidyl transferase dUTP nick end labeling (TUNEL) assay was performed using an *in situ* cell death detection kit conjugated with fluorescein isothiocyanate (Roche Diagnostics GmbH, Mannheim, Germany, 11684795910). Retinal tissue sections from WT, *Rho*^ΔI255/+^, and *Rho*^ΔI255/ΔI255^ mice at PN11, PN13, PN15, PN18, PN20, PN22, PN24, PN26, PN30, PN40 and PN60 were collected for TUNEL staining. DAPI (D9542, Sigma) was used as a nuclear counterstain.

### Calpain *in situ* activity assay

Calpain activity was investigated with an enzymatic *in situ* activity assay [51]. Briefly, unfixed retinal cryosections at 9 distinct time points (PN11, PN13, PN15, PN18, PN20, PN24, PN30, PN40, and PN60) were dried at 37°C for 15 min, and rehydrated for 15 min in calpain reaction buffer (CRB; 25 mM HEPES, 65 mM KCl, 2 mM MgCl_2_, 1.5 mM CaCl_2_ in ddH_2_O) and then incubated at 37°C for 3.5 h in the dark in CRB with 2 mM dithiothreitol (DTT) and t-BOC-Leu-Met-CMAC (25 µM; A6520, Thermo Fisher Scientific, OR, USA). Retinal sections were washed with PBS and incubated with ToPro3 (1:1000 in PBS, Thermo Fisher Scientific, OR, USA) at RT for 25 min, which was used as the nuclear counterstain. Afterward, sections were washed with PBS and mounted using Vectashield without DAPI (Vector Laboratories Inc., Burlingame, CA, USA) for immediate visualization.

### PARP *in situ* activity assay

This assay allows for resolving the overall activity of poly(ADP-ribose)polymerase (PARP) *in situ* on unfixed tissue sections [94]. Retinal sections from the aforementioned 9 time points were dried at 37°C for 15 min and rehydrated for 10 min in Tris buffer. The PARP reaction mixture containing 1 mM DTT, 50 μM 6-Fluo-10-NAD^+^ (N 023, Biolog, Bremen Germany), and PARP buffer (10 mM MgCl_2_ in 100 mM Tris buffer with 0.2% Triton X100) was applied to the sections for 3.5 h at 37 °C in the dark. After 3 times washing with PBS, the sections were mounted with DAPI (Vectashield with DAPI; Vector Laboratories, Burlingame, CA, USA) for immediate visualization.

### Transmission electron microscopy

WT and *Rho*^ΔI255/+^ mice eyecups were fixed at PN20 with 2.5% glutaraldehyde, 2 % PFA, and 0.1 M sodium cacodylate buffer (pH 7.4, Electron Microscopy Sciences, Germany) overnight at 4°C. After rinsing, samples were postfixed in 1 % OsO4 for 1.5 hours at room temperature, washed in cacodylate buffer, and dehydrated with 50 % ethanol. Tissues were counterstained with 6 % uranyl acetate dissolved in 70 % ethanol (Serva, Germany), followed by graded ethanol concentrations up to 100 % and Propylenoxide. The dehydrated samples were incubated in 2:1, 1:1, and 1:2 mixtures of propylene oxide and Epon resin (Serva, Germany) for 1 hour each. Finally, samples were infiltrated with pure Epon for 2 hours. Samples were embedded in fresh resin in block molds and cured 3 days at 60 °C. Ultrathin sections (50 nm) were cut on a Reichert Ultracut S (Leica, Germany), collected on copper grids, and counterstained with Reynold’s lead citrate. Sections were analyzed with a Zeiss EM 900 transmission electron microscope (Zeiss, Jena; Germany) equipped with a 2k x 2k CCD camera. During this experiment, we focused our investigation on the inferior retina because the degeneration in the inferior retina is more severe than that of the superior retina.

### Post-embedding immunogold staining (IGS)

Retinal samples were fixed in 0.05% glutaraldehyde and 4% paraformaldehyde in 0.1 M phosphate buffer saline (PBS), pH 7.2 (Electron Microscopy Sciences, Germany), for 4h at 4°C and further sliced to yield small sections. After rinsing in PBS, sections were dehydrated through an ethanol series (15 min in each of 50%, 70%, 95%, 2×100% ethanol) on ice, and infiltrated with LR-White resin (Biotrend, Germany; 20 min in each of 2:1, 1:1, and 1:2 ethanol:LR-White). After two changes of LR-White, tissues were infiltrated in fresh LR-White for 24 h at 4° C. Finally, samples were placed in flat embedding molds containing fresh LR-White and polymerized for 24 h at 56°C.

Ultrathin sections (70 nm thickness) were cut on a Reichert Ultramicrotome S (Leica, Germany) and collected on Formvar-coated nickel grids for immunogold staining. Grids were washed with 0.5 M glycine in PBS and then blocked in 2.5% goat serum, 2.5% ovalbumin, 0,1% cold water fish skin gelatin and 0,1% bovine serum albumin in PBS containing 0,1% Triton X-100 (Biotrend, Germany) for 30 min at room temperature before incubation with mouse anti rhodopsin primary antibody (Table S1) diluted 1:30 in blocking buffer over night at 4 C°. As a negative control, the corresponding grid for each sample was included in parallel without the primary antibody incubation. Next, the grids were washed the grids were washed in PBS 6 times in PBS, and labeled with 12 nm colloidal gold conjugated goat anti-mouse immunoglobulins (Jackson ImmunoResearch Laboratories) diluted 1:20 in the above blocking buffer but without goat serum and ovalbumin for 1 h at R.T. Grids were then washed with PBS (6×5 min each), postfixed in 1% glutaraldehyde (Electron Microscopy Sciences, Germany) in PBS and rinsed with distilled water. Sections were counterstained with 3% aqueous uranyl acetate (Serva, Germany) for 3 min, followed by a final rinse with distilled water and air-dried. Sections were analyzed with a Zeiss EM 900 transmission electron microscope (Zeiss, Jena; Germany) equipped with a 2k x2k CCD camera.

### Electroretinography (ERG)

Mice were dark-adapted overnight. Subsequently, they were anesthetized with a subcutaneous injection of ketamine (66.7 mg/kg of body weight) mixed with xylazine (11.7 mg/kg of body weight) and diluted in 0.9% NaCl saline. Tropicamide drops (Pharma Stulln, Stulln, Germany) were applied to each eye for pupil dilation. All handling procedures were performed under dim red light, and recordings were conducted under dark conditions. The animals were positioned in the prone position on a temperature-controlled mat and gold-wire ring electrodes, moisturized with methylcellulose (OmniVision GmbH, Puchheim, Germany), contacted the surface of both corneas for binocular recordings. Two short stainless-steel needle electrodes (Sei Emg s.r.l., Cittadella, Italy) were used as the reference and ground electrodes. Full-field ERG recordings were performed with the Espion E^3^ console connected to a computer, a 32-bit amplifier, and a Ganzfeld Bowl (Diagnosys, LLC, Lowell, MA, USA). Amplifier cutoff frequencies were 0.3 Hz (lower) and 300 Hz (upper). Initially, a dark-adapted (scotopic) single-flash ERG series, ranging from 1 mcd*s/m^2^ to 30 cd*s/m^2^, was recorded. Following 5 min light adaptation with a rod-saturating background of 30 cd*s/m^2^, a light-adapted (photopic) single-flash ERG series, ranging from 10 mcd*s/m^2^ to 30 cd*s/m^2^, was recorded on the same background. Each recording step was averaged 10–15 times, with inter-stimulus intervals of 2 s (1–300 mcd*s/m^2^), 5 s (1–3 cd*s/m^2^), or 10 s (10–30 cd*s/m^2^). B-wave amplitudes were measured from the trough of the a-wave to the peak of the b-wave. Box-and-whisker plots were produced with Microsoft Excel (Microsoft Corp., Redmond, WA, USA), and figures were prepared using the CorelDRAW X5 software (Corel Corporation, Ottawa, ON, Canada).

### Scanning-laser ophthalmoscopy (SLO) and optical coherence tomography (OCT)

*In vivo* imaging was performed as previously described [95, 96]. Animals were examined directly after the ERG session with continuing anesthesia. A 100 dpt contact lens was placed on the cornea after applying hydroxypropyl methylcellulose to avoid dehydration. SLO imaging was performed together with OCT on the Spectralis™ HRA+OCT device featuring a superluminescent diode at 870 nm as low coherence light source using the proprietary software package Eye Explorer version 5.3.3.0 (Heidelberg Engineering, Heidelberg, Germany). Each two-dimensional B-Scan recorded at 30° field of view contains up to 1536 A-Scans, acquired at 40,000 scans per second. Data was exported as 8-bit greyscale images. For layer quantification, image data was processed with the ImageJ software package (http://rsb.info.nih.gov/ij/), and reflectivity profiles between the ganglion cell layer and the retinal pigmented epithelium were extracted from the OCT scan as described previously [97]. To determine the thickness of the inner/outer retina, the size of GCL, IPL & INL / OPL, ONL & IS+OS was added, respectively.

### Microscopy and image analysis

All immunostaining samples were analyzed using a Zeiss Axio Imager Z1 ApoTome microscope, AxioCam MRm camera, and Zeiss Zen 2.3 software in Z-stack at 10x, 20x, or 40x magnification. TUNEL-positive cells in the ONL of at least three sections per group were manually counted for quantitative analysis. The percentage of TUNEL-positive cells was calculated by dividing the number of positive cells by the total number of ONL cells. Photoreceptor cell rows were assessed by counting the individual nuclei rows in one ONL and averaging the counts. The intensity mean value of RHO immunofluorescence was measured by contouring the ONL and IS areas, and the intensity average of GFAP immunofluorescence was measured by outlining the entire retina layer from the GCL to the OS, using Zeiss Zen software. To correct for differences in background intensity, an area devoid of GFAP staining was used, and the average intensity in this area was subtracted from the average intensity of GFAP, *i.e*., the relative MFI of GFAP protein. OS thickness was measured via the distance tool in Zeiss Zen software.

For wholemount retina, cells stained for S-opsin and M-opsin in superior and inferior regions from WT and *Rho*^ΔI255/+^ retinae were counted by Image J software (http://imagej.nih.gov/ij). We selected an area of 30,000 µm^2^ area in the corresponding region of the retina for cell counting analysis. In Image J, the plugin ITCN automatic cell counting analyzer was used to count cells after inverting images and converting them to 8-bit. At least 3 wholemount retinae (n=3) from different animals were included. Graphs were prepared in GraphPad Prism 10.2.1 for Windows; Inkscape software was used for image processing.

### Statistics

Statistical analysis was performed using GraphPad Prism 10.2.1 Software. A two-way ANOVA was used to evaluate differences in the percentage of TUNEL positive cells in the ONL, ONL rows, cone numbers per 100 µm of the retina, the mean number of microglial cells in the ONL and the subretinal space, the percentage of caspase-3, PARP, and calpain positive cells in the ONL. A one-way ANOVA compared the RHO mean fluorescence intensity and OS thickness between WT, *Rho*^ΔI255/+^, and *Rho*^ΔI255/ΔI255^ mice. When a 0.05 level of significance was found, Bonferroni post-hoc tests were performed to compare mutant groups with WT. An unpaired two-tailed *t*-test was used to evaluate the differences between S-cone and M-cone numbers and the relative MFI of GFAP protein in the retina. Quantitative data are shown as the mean ± standard deviation (SD). *p* < 0.05 was considered statistically significant.

## Author contributions

Bowen Cao: Data curation, Formal analysis, Investigation, Validation, Visualization, Writing – original draft, Writing – review & editing; Yu Zhu: Investigation, Validation, Visualization, Writing – original draft, Writing – review & editing; Alexander Günter: Data curation, Formal analysis, Investigation, Visualization, Writing – original draft; Ellen Kilger: Visualization, Writing – review & editing; Sylvia Bolz: Investigation, Methodology, Writing – review & editing; Christine Henes: Investigation; Regine Mühlfriedel, Data curation, Formal analysis, Resources, Validation, Visualization, Writing – original draft; Mathias W. Seeliger: Formal analysis, Methodology, Resources, Validation, Visualization, Writing – original draft, Writing – review & editing, François Paquet-Durand: Conceptualization, Funding acquisition, Investigation, Methodology, Resources, Validation, Writing – review & editing; Blanca Arango-Gonzalez: Conceptualization, Data curation, Funding acquisition, Investigation, Methodology, Resources, Supervision, Validation, Visualization, Writing – review & editing; Marius Ueffing: Conceptualization, Funding acquisition, Investigation, Resources, Supervision, Validation, Visualization, Writing – review & editing.

## Funding statement

This research was supported by grants from the Tistou and Charlotte Kerstan Foundation (RHO-Cure program, to Marius Ueffing), URL: http://www.kerstanstiftung.org, Fighting Blindness Canada (FBC, to Marius Ueffing and Blanca Arango-Gonzalez), URL: Transformative Research Award: - TargetVCP: https://www.fightingblindness.ca), the Zinke Heritage foundation (to Marius Ueffing), and the ProRetina foundation (to Marius Ueffing), URL: www.pro-retina-stiftung.de.

## Acknowledgements

We would like to thank Anne-Sophie Petremann-Dumé and Norman Rieger for excellent technical assistance.

## Conflict of interest statement

All authors of this manuscript acknowledge the PLOS Genetics policy on conflicts of interest and none of the authors of this manuscript declares a conflict of interests.

---

**Supplementary Figure. 1.**
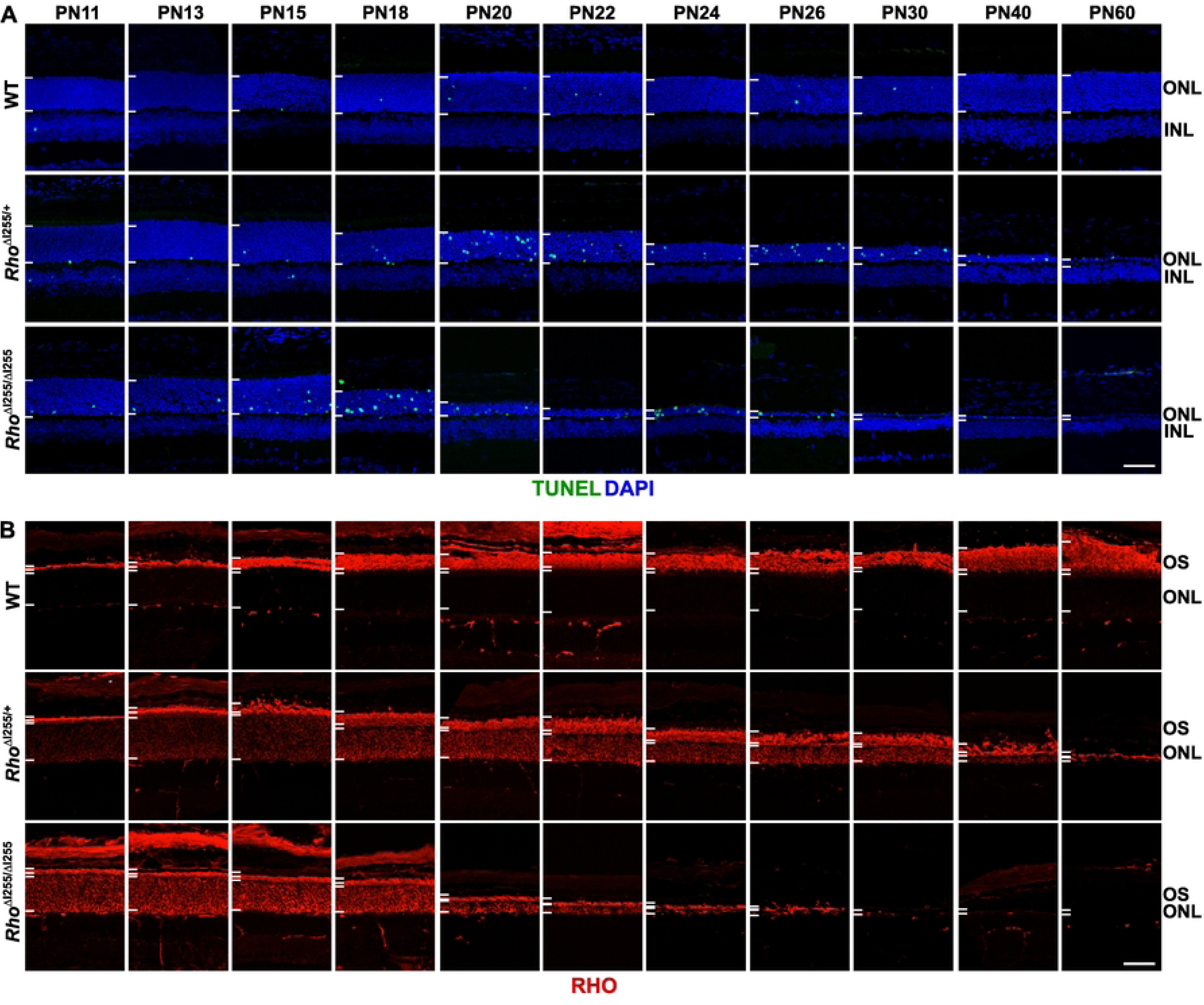

**Supplementary Figure. 2.**
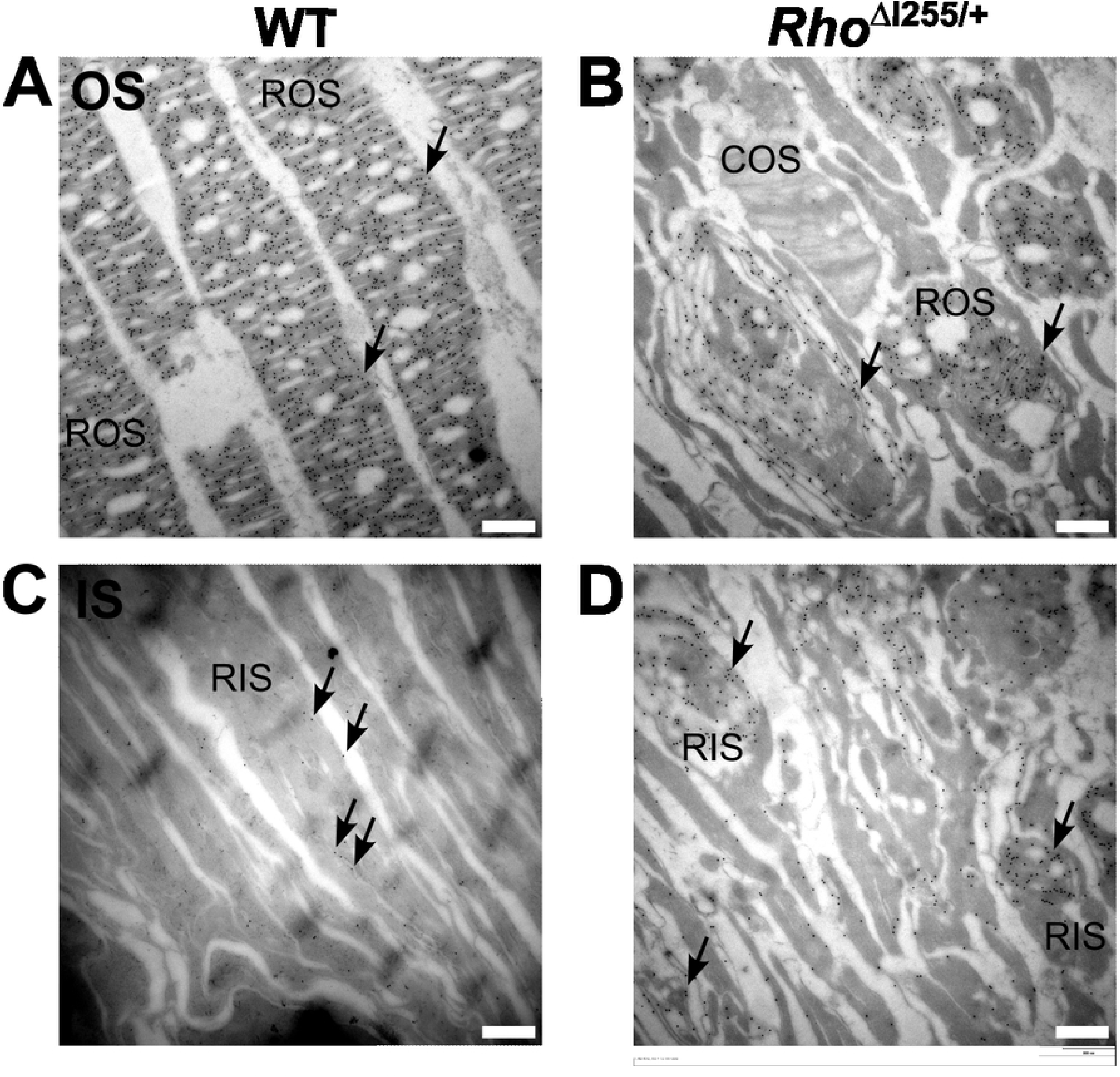

**Supplementary Figure. 3.**
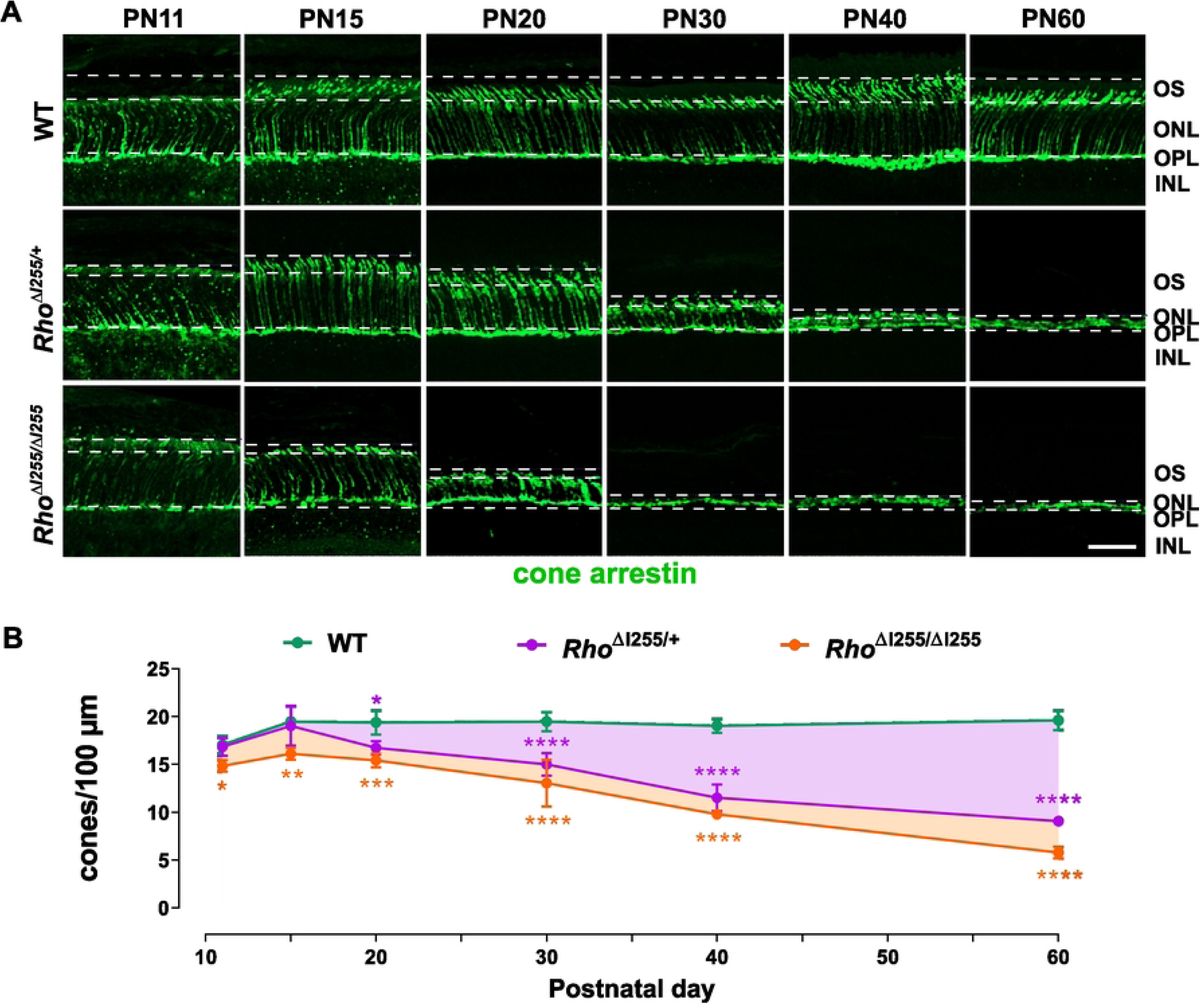

**Supplementary Figure. 4.**
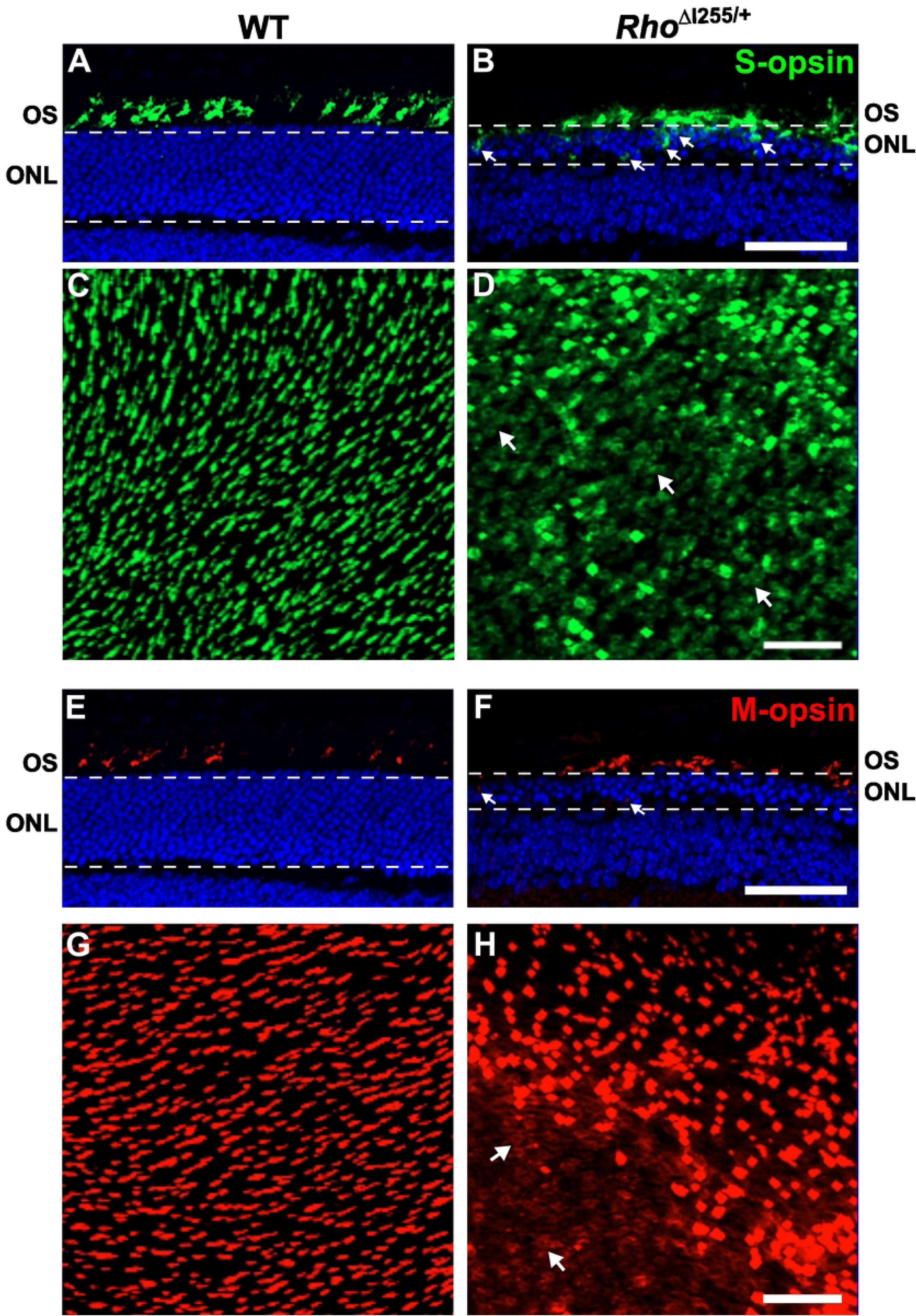

**Supplementary Figure. 5.**
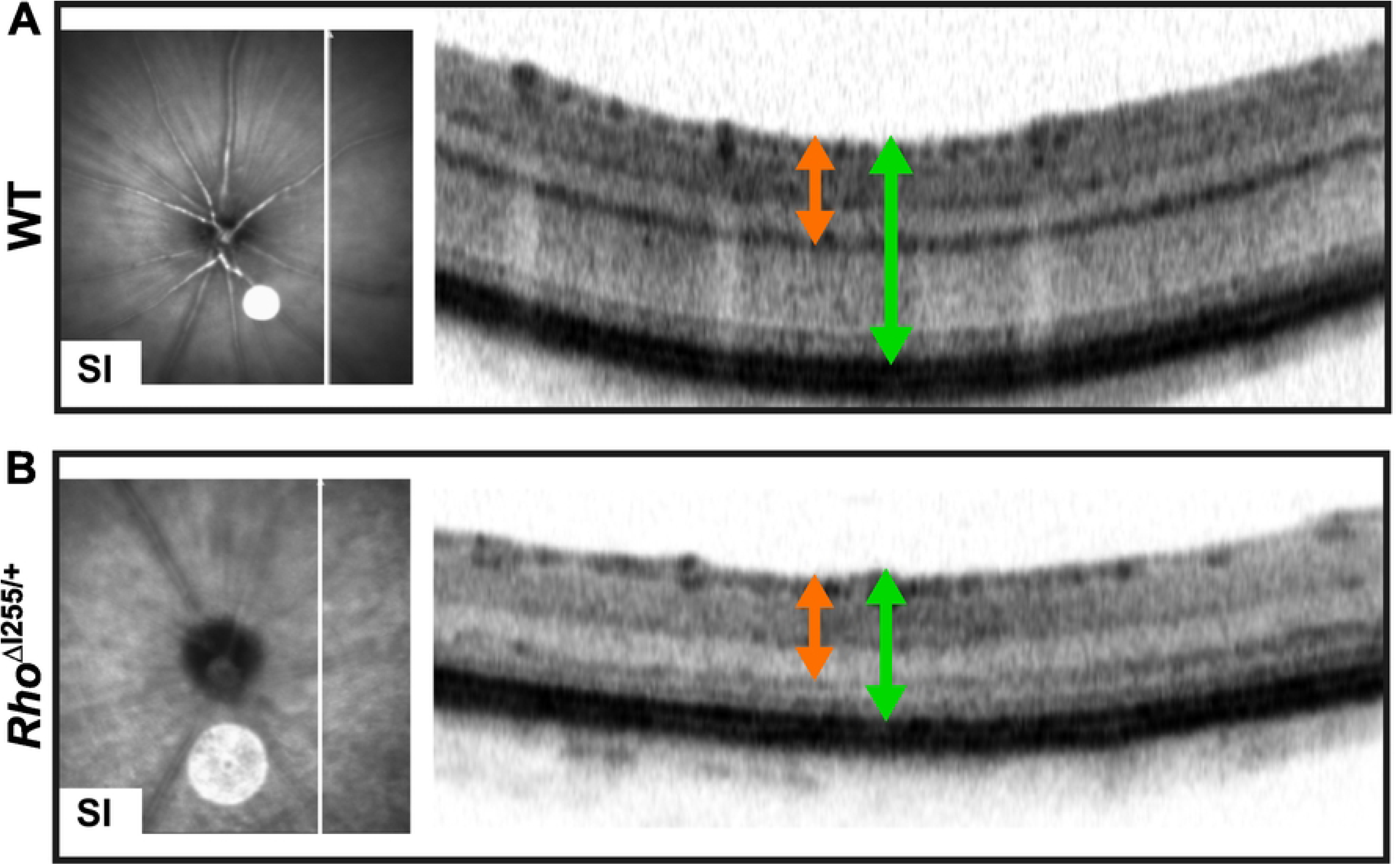

## Notes

### Competing Interest Statement

The authors have declared no competing interest.

